# Reproductive and cognitive effects in carriers of recessive pathogenic variants

**DOI:** 10.1101/2024.09.30.615774

**Authors:** Hila Fridman, Gelana Khazeeva, Ephrat Levy-Lahad, Christian Gilissen, Han G. Brunner

## Abstract

The genetic landscape of human Mendelian diseases is shaped by mutation and selection. Selection is mediated by phenotypic effects which interfere with health and reproductive success. Although selection on heterozygotes is well-established in autosomal dominant disorders, convincing evidence for selection in carriers of pathogenic variants associated with recessive conditions is limited, with only a few specific cases documented.

We studied heterozygous pathogenic variants in 1,929 genes, which cause recessive diseases when bi-allelic, in a cohort of 378,751 unrelated European individuals from the UK Biobank^1^. We assessed the impact of these pathogenic variants on reproductive success. We find evidence for fitness effects in heterozygous carriers for recessive genes, especially for variants in constrained genes across a broad range of diseases. Our data suggest reproductive effects at the population level, and hence natural selection, for autosomal recessive disease variants. We further show that variants in genes that underlie intellectual disability are associated with reduced cognition measures in carriers. In concordance with this, we observe an altered genetic landscape, characterized by a threefold reduction in the calculated frequency of biallelic intellectual disability in the population relative to other recessive disorders. The existence of phenotypic and selective effects of pathogenic variants in constrained recessive genes is consistent with a gradient of heterozygote effects, rather than a strict dominant-recessive dichotomy^2^.

## Main

Together with mutation, selection is the driving force behind genetic diversity, as it constantly changes the genetic landscape. For autosomal-dominant conditions, such selection is often immediate and strong. An example is severe intellectual disability (ID), where new mutations predominate because affected individuals rarely reproduce^3,4^. Similarly, some de novo copy number variants (CNVs) are under strong purifying selection, while intermediate selection is observed for inherited CNVs with reduced penetrance^5^. For recessive conditions, there is a strong selection against biallelic pathogenic variants resulting in severe disease. Recent work suggests that milder phenotypes may be more frequent in heterozygotes than previously assumed^6^, suggesting that the traditional dominant-recessive dichotomy should be replaced with a graded system of variant effects^2^. Simulations show that, because heterozygotes for recessive disease variants are much more common in the population than individuals with biallelic pathogenic variants, their combined selective effects could be substantial^7-9^. In specific instances, a positive selective effect in heterozygotes has been proposed for some recessive conditions that have high carrier rates, such as Phenylketonuria (PKU) and Cystic fibrosis (CF)^10-12^. Evidence for positive selection for these conditions was mainly correlative and based on relatively high population frequencies. How much heterozygote selection impacts the recessive disease landscape is currently not known. Nonetheless, if there is substantial negative selection on heterozygous pathogenic alleles, this could affect the frequency with which recessive diseases occur.

### Heterozygote selection in recessive genes

To evaluate negative selection against heterozygotes for pathogenic and likely pathogenic variants (PLPs) in recessive disease genes, we used exome sequencing data from 378,751 unrelated European individuals of the UK Biobank, and identified 54,758 different presumable PLPs in a curated set of 1,929 autosomal-recessive (AR) disease genes (**Fig. 1a, Methods, Supplementary Table 1**). Briefly, most PLPs (80.9%: 44,277/54,758) were selected based on loss of function (LoF) mechanism and rarity (population frequency <1%). PLPs were also collected from ClinVar and the Dutch diagnostic laboratories database VKGL (14.9%: 8,208/54,758). Missense variants that were assessed as PLPs by at least 2/3 different databases were also considered as PLPs (4.1%: 2,273/54,758) (**Methods**). We found that on average, each individual is heterozygous for ∼1.9 AR PLPs (**Figure 1b**). Most of these PLPs (99.9%) are rare, with a heterozygote frequency of 0-0.5% in the cohort (**Supplementary data Fig. 1a**,**b**). These estimates for heterozygosity in the UK Biobank are very similar to those that were previously obtained in Dutch and Estonian populations^9^. In line with the expected deleteriousness of the PLPs, we observe a depletion of more common PLPs in highly constrained genes (**Fig. 1c**). For our analyses, we selected PLPs that were absent in homozygous state and observed in heterozygous state in no more than 20 individuals within our cohort (92.3%: 50,568/54,758). We additionally excluded 2,143 individuals with PLPs in a potential biallelic state, leaving a final set of 376,608 individuals.

**Fig. 1.**
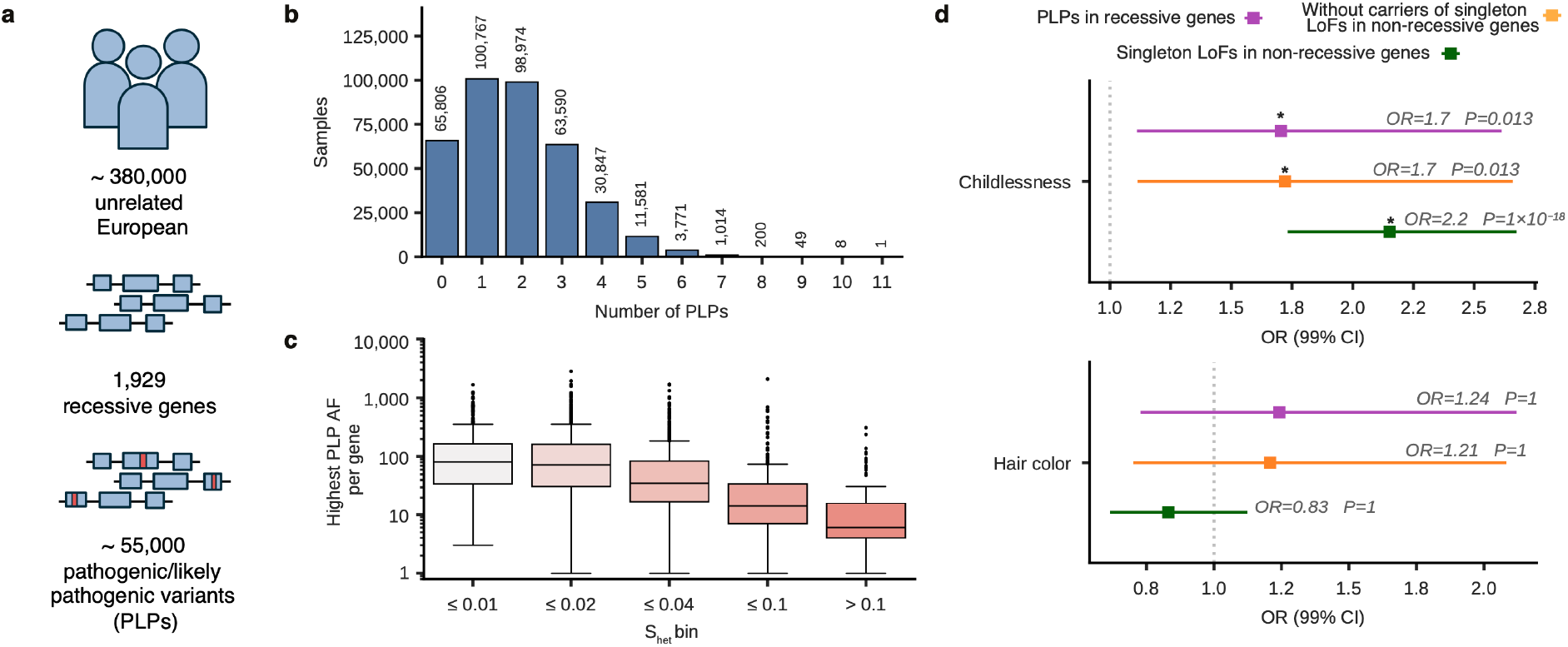
Association of genetic burden for recessive disease with childlessness. **a**, Overview of the study: selection of individuals, recessive genes and pathogenic variants. **b**, Distribution of the number of PLPs seen per individual. **c**, Distribution of the highest AF of PLPs per gene for different selection constraint (s_het_) bins. **d**, Effect of genetic burden on childlessness and hair color (as a control phenotype) for heterozygous PLPs in recessive genes (purple), heterozygotes for PLPs in recessive genes excluding carriers of LoF in non-recessive genes (orange) and singleton LoFs in all other constrained (s_het_ > 0.15) non-recessive genes (green). Colored lines indicate odds ratios for the phenotypes with 99% confidence intervals; dashed gray line indicates odds ratio of 1; p-values are adjusted for multiple testing using Bonferroni correction; significant associations are marked with an asterisk.

Reduced reproductive success has recently been associated with damaging genetic variation in constrained genes throughout the human genome^13^. To determine whether increased childlessness is also detectable for carriers of damaging variants in a curated subset of 1,929 autosomal recessive genes^9^ we took a similar approach (**Methods**). For each individual in the UK Biobank we calculated “genetic burden”, as previously described^13^ (**Methods**), based on the combined s_het_ scores from Seplyarskiy et al.^14^ for all 1,929 AR disease genes in which that individual harbored PLPs. We evaluated the association between genetic burden and childlessness using logistic regression while correcting for age and the first 40 genetic principal components. Bonferroni correction was applied to all p-values to account for multiple testing (**Methods**).

Logistic regression showed a positive association of genetic burden in 1,929 AR disease genes with “childlessness” (odds ratio: 1.7, p-value: 0.013) (**Fig. 1d**). Results were very similar when repeating the analysis for three other constraint scores^13,16^ (**Supplementary data Fig. 2, Supplementary Table 2**). As a control, we performed the same association analysis for “hair color”, and found no significant association (odds ratio: 1.24, p-value: 1) (**Fig. 1d, Supplementary Table 2**). The analysis of the association between childlessness and carriership status for PLPs in recessive genes did not reveal any significant findings (**Supplementary data Fig. 3, Supplementary Table 3**).

To confirm that these results were not confounded by the effect of PLPs in other, non-recessive genes, we reanalyzed our data after removing 11,557 individuals heterozygous for LoF singleton variants in any highly constrained non-recessive gene (n=1,908, defined as Weghorn et al.^15^ s_het_ ≥ 0.15). We also repeated the analysis incorporating heterozygote status for these genes as a covariate and obtained very similar results in both analyses (**Fig. 1d, Supplementary Table 4, Methods**). This shows that the effect of heterozygous PLPs in constrained recessive genes is independent of the effect of LoF singletons in other constrained genes. We note that the odds ratios for the association of heterozygous PLPs with childlessness do not differ significantly between 1,929 recessive and 1,908 non-recessive highly constrained genes (Wald statistic: 1.4, two-sided p-value: 0.236; **Methods**), even though the significance is much stronger for non-recessive genes (**Supplementary data Fig. 4, Supplementary Table 5)**.

### Heterozygote cognitive effect in AR-ID genes

Having found a correlation between the genetic burden in heterozygous carriers of recessive diseases and childlessness, we asked whether this correlation could be partly mediated by factors such as fertility, disease, deprivation, or reduced cognitive abilities. There is abundant evidence from previous work that cognitive and behavioral factors are important in involuntary childlessness. For example, in a cohort of full siblings, the strongest association with childlessness was observed for mild intellectual disability in both males and females^16^. In addition, a study on the UK Biobank found that the genetic burden of damaging heterozygous variation across the entire genome was associated with markedly reduced reproductive success, primarily owing to increased childlessness. The genetic burden of damaging genetic variation correlated negatively with cognitive and behavioral traits in both sexes^13^.

To test these hypotheses we performed generalized linear regressions with the genetic burden in the 1,929 AR gene set as predictor for the number of ICD-10 diagnoses (n=376,608) as a proxy for health status, for ICD-10 codes associated with infertility (n=889), and for educational attainment (in years of education, n=373,090) as proxy for cognitive ability. We found that individuals with a higher genetic burden have higher numbers of ICD-10 codes (relative change: 40%, p-value: 5.73×10^-6^) and lower educational attainment (effect size: -1.86, p-value: 2.65×10^-7^) (**Fig. 2a, Supplementary Table 2, Supplementary data Fig. 5**). The association of genetic burden with childlessness was not explained by the number of ICD-10 diagnoses, infertility (**Supplementary data Fig. 6**) or any measures of deprivation (**Supplementary data Fig. 7a**). As another measure of cognition, we assessed an individual’s fluid intelligence score (n=120,759), which estimates the capacity to solve problems that require logic and reasoning ability (**Methods**). Fluid intelligence did not reach significance, likely because of reduced power for this variable due to the smaller number of samples (**Supplementary data Fig. 5**). Similarly, there was no significant correlation of genetic burden with infertility (**Supplementary Table 2**). As an additional control we repeated the regressions using rare synonymous variants in recessive genes (**Methods)**. After 300 resamplings, the association strength distributions for synonymous variants did not overlap with those of PLPs, except for hair color and infertility (**Fig. 2b**).

**Fig. 2.**
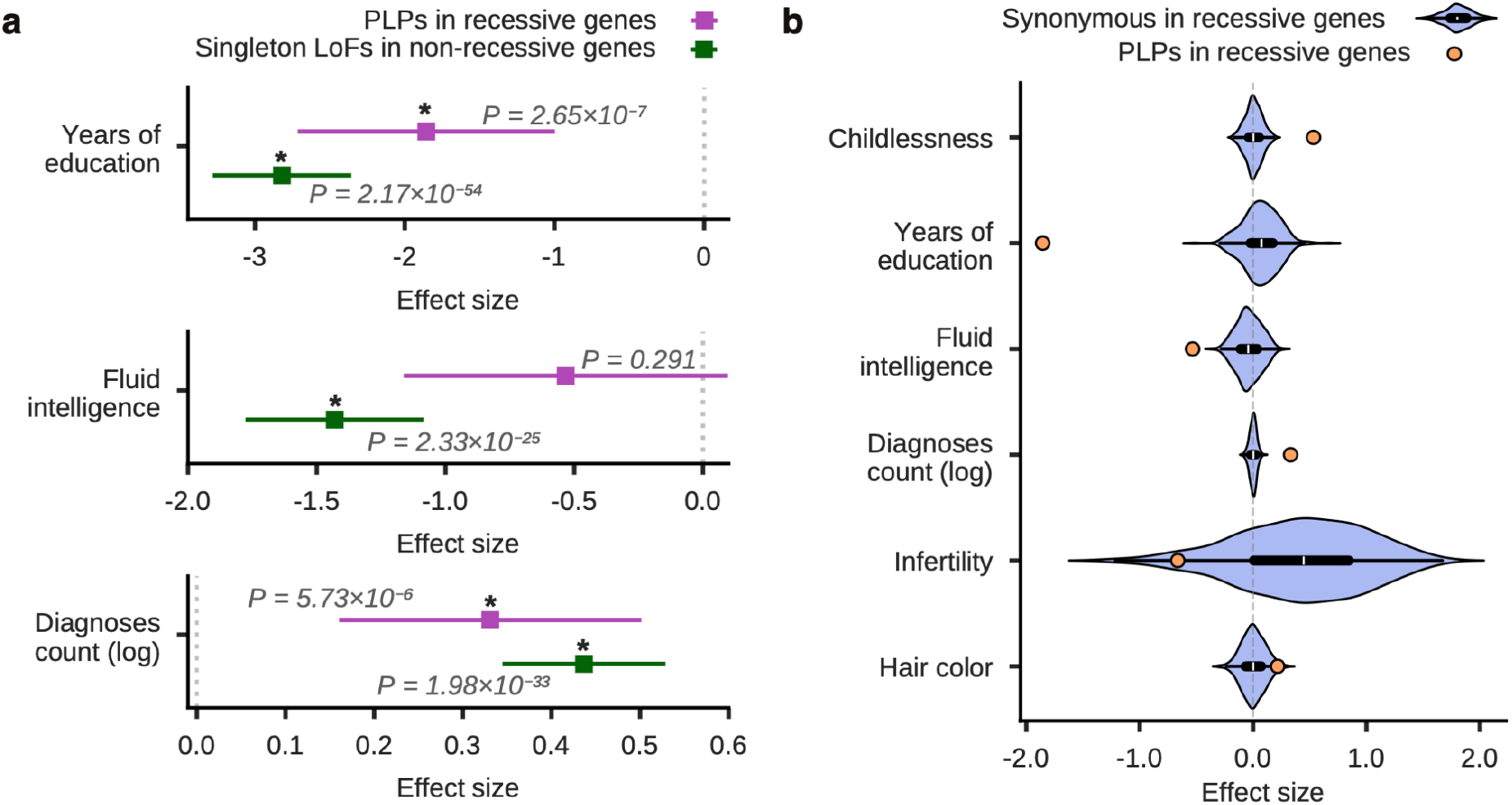
Association of recessive genetic burden with cognitive and disease-related phenotypes. **a**, Effect of genetic burden on educational attainment (measured in years of education), fluid intelligence score and log-transformed number of ICD-10 diagnoses for PLPs in all recessive (purple) and singleton LoFs in non-recessive highly constrained (green) genes. Colored lines indicate effect sizes (estimated regression coefficients) with 99% confidence intervals; dashed gray line indicates the effect size of 0; p-values are adjusted for multiple testing using Bonferroni correction; significant associations are marked with an asterisk. **b**, Distribution of the estimated effect sizes (regression coefficients) for synonymous variants in recessive genes re-sampled 300 times (light purple violin plots) compared with the estimated effect size for PLPs in recessive genes (orange dots).

Given the evidence for possible mediation of childlessness through cognitive abilities, we examined whether the observed effect of genetic burden on childlessness is equally distributed across 13 non-overlapping disorder groups, including ID. After multiple testing correction only recessive ID genes showed a significant negative association with childlessness (odds ratio: 2.76, p-value: 0.024), educational attainment (effect size: -3.93, p-value: 2.84×10^-8^) and log-transformed number of ICD-10 diagnoses (effect size: 0.55, p-value: 3.79×10^-4^) (**Fig. 3a, Supplementary Table 6-9**). The association with childlessness for ID genes does not change when controlling for measures of deprivation (**Supplementary data Fig. 7b**).

**Fig. 3.**
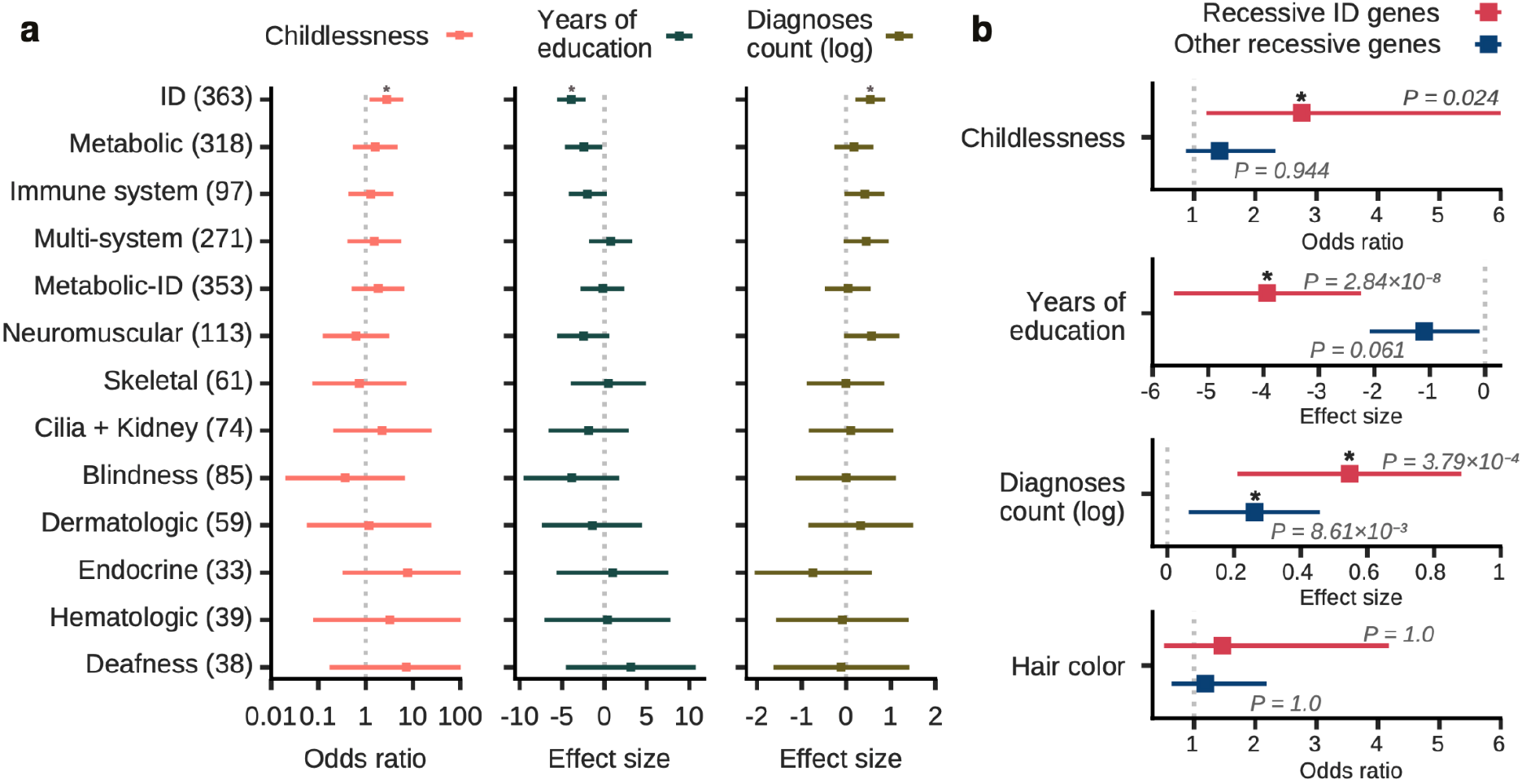
Association of childlessness, educational attainment, log-transformed number of ICD-10 diagnoses for different disorder groups. **a**, Effect of genetic burden on childlessness (peach), educational attainment (measured in years of education; green) and log-transformed number of ICD-10 diagnoses (brown) for PLPs in recessive genes divided into 13 non-overlapping disorder groups. Colored lines indicate odds ratio (for childlessness) or effect sizes (estimated regression coefficients) with 99% confidence intervals; dashed gray line indicates the odds ratio of 1 (for childlessness) or the effect size of 0; p-values are adjusted for multiple testing using Bonferroni correction; significant associations are marked with an asterisk. **b**, The comparison between PLPs in recessive ID genes (red) and all other recessive genes (blue) for the effects of genetic burden on childlessness, educational attainment (measured in years of education), log-transformed number of ICD-10 diagnoses and hair color (as a control phenotype). Colored lines indicate odds ratio (for childlessness and hair color) or effect sizes (estimated regression coefficients) with 99% confidence intervals; dashed gray line indicates the odds ratio of 1 (for childlessness and hair color) or the effect size of 0; p-values are corrected for multiple testing; significant associations are marked with an asterisk.

The estimates for these associations were more pronounced for ID genes compared to other recessive genes (**Fig. 3b, Supplementary data Fig. 8**), with a significant difference between gene groups detected only for educational attainment (Wald statistic: 14.8, p-value: 0.0001). The associations of recessive genetic burden with childlessness, years of education and number of ICD-10 diagnoses are more pronounced for both ID and other recessive genes in comparison with synonymous variants (**Supplementary data Fig. 9**). As genetic burden is dependent on cumulative s_het_ scores per individual, we explored whether recessive ID is enriched for genes with high s_het_ scores (**Supplementary data Fig. 10**). We found that the median s_het_ score of ID genes is indeed significantly higher than the median s_het_ score of recessive genes for the other 12 disorder groups (Mood’s median test statistic: 13.3, p-value: 0.0003). To investigate the possibility that the effects observed for ID genes are solely due to the enrichment for highly constrained genes, we constructed simulated gene sets from recessive non-ID genes with the same s_het_ distribution as ID genes with 20 resamplings (**Methods; Supplementary data Fig. 11**). We find that the distribution of association strength for simulated gene sets with similar selection profiles does not overlap with those estimated for recessive ID genes in case of childlessness, educational attainment and fluid intelligence score (**Supplementary data Fig. 12**). Thus, we find that the burden of pathogenic variants in a wide range of recessive genes correlates with increased childlessness, but that the impact on reduced cognition is stronger for ID genes.

### Analyses in males and females

The causes of childlessness in men and women only partly overlap. For instance, a Finnish-Swedish study on early childhood diseases in full siblings found a broadly similar pattern in men and women with some notable differences for endocrine disorders, as would be expected^16^. Such differences are likely at least partly due to underlying genetic differences. A GWAS for genes influencing reproductive behavior found few differences between the sexes^17^, but a study of polygenic risk scores of 9,942 Swedish twin individuals was suggestive of substantial genetic effects, although these were mediated by different gene sets in men and women^18^. A study of a different cohort of 7,820 twin individuals from Finland documented educational differences in completed fertility. This study found that poorly educated men, and highly educated women, are least likely to have any children, and that these groups have lower completed fertility in general. Behavioral genetics analysis of this sample further suggested that the association between education and having any children in both sexes is influenced by factors shared by co-twins and that these factors are genetic rather than environmental^19^.

Given this previous evidence^13,18,19^, we examined whether the effects of a higher recessive genetic burden differed between males and females (**Supplementary data Fig. 13; Supplementary Tables 10-11**). After including the interaction term between genetic burden and sex (**Methods**) we found no significant differences between males (n=174,470) and females (n=202,138) for childlessness (p-value: 0.293) and educational attainment (p-value: 0.494).

### The genetic landscape

We hypothesized that the selective effect seen for PLPs in recessive ID genes could be reflected in the frequency and distribution of these variants in the population. We refer to this as the “genetic landscape”. Specifically, we hypothesized that, based on our previous results, the genetic landscape of ID genes would include fewer high-frequency PLPs than other recessive genes. Following methodology from Fridman et. al^9^, we measured this effect by the consanguinity ratio (CR) which is the ratio between the predicted frequency of couples at-risk for an affected child for first-cousin couples to that for random couples (**Methods**). In the UK Biobank cohort, we found that the CR is significantly elevated only for recessive ID genes, with no such elevation observed for any of the other disorder groups (permutation test, p-value after Bonferroni correction=0.005, **Methods**) (**Fig. 4a, Supplementary data Fig. 14, Supplementary Table 12**). In fact, if ID had the same variant distribution as other recessive diseases, its incidence would be threefold higher (CR=67 for ID genes, versus CR=19 for recessive genes when excluding ID) (**Fig. 4a)**. The same landscape was previously observed in smaller Dutch and Estonian cohorts^9^ (**Supplementary Table 12**).

**Fig. 4.**
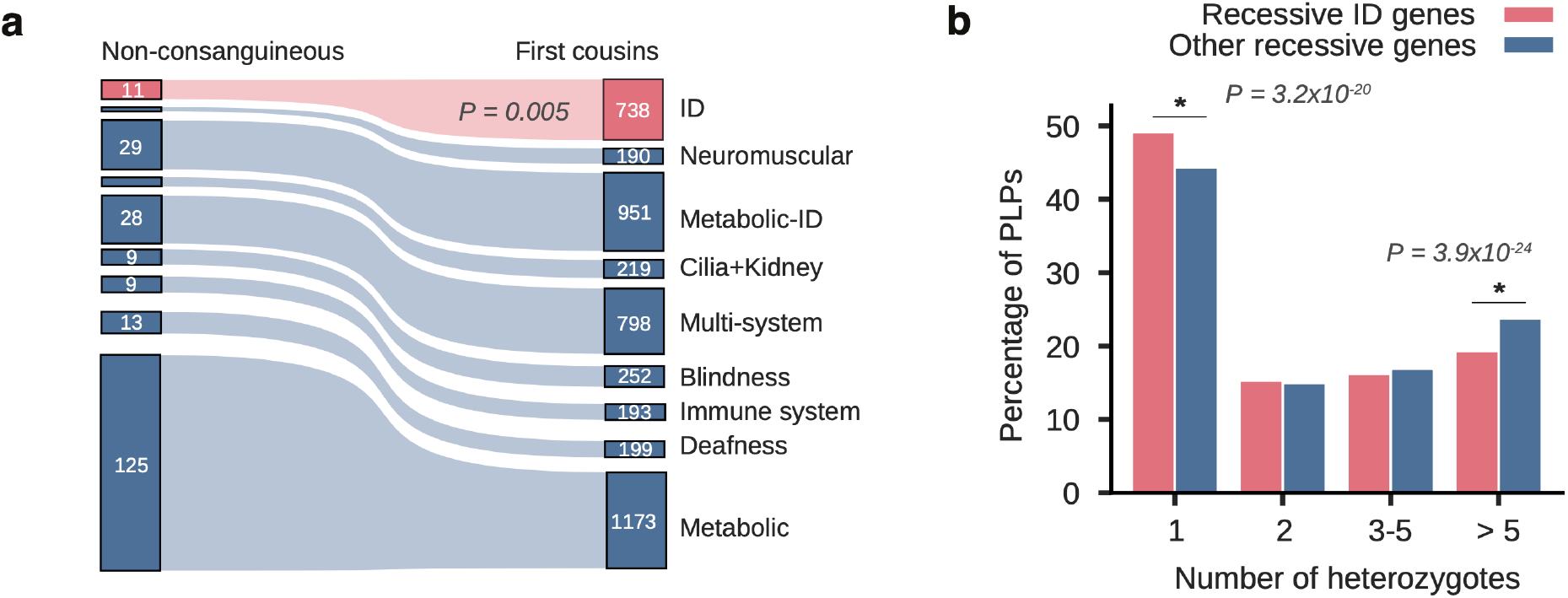
Consanguinity effects and the genetic architecture for various disorder groups. **a**, The expected number of affected offspring for different disorder groups, calculated per 100,000 births, in non-consanguineous and first cousins matings. Recessive ID genes are in red and all other recessive genes are in blue. **b**, The distribution of the number of heterozygotes for PLPs in recessive ID genes (red) and in all of the other recessive genes (blue).

CR reflects the genetic landscape of different disorder groups in multiple ways that involve selection and is relevant to the occurrence of disease. These include the fraction of genes with higher s_het_ scores, depletion of high-frequency alleles, and the distribution of PLP allele frequencies. All of these factors correlate with the CR score as expected (**Fig. 4b, Supplementary Table 13**). While these factors likely contribute to the landscape that we observe, they are certainly not a complete explanation. A complex interplay of positive, neutral, and negative selection, population history, environmental changes, genetic drift, and various other factors have all contributed over time, and probably still contribute today.

We observed that the average PLP allele frequency (AF) per disorder group showed a high correlation between the UK Biobank and Dutch cohorts (Pearson correlation coefficient: 0.79, p-value: 0.001) (**Supplementary data Fig. 15**). This correlation demonstrates strong similarity of the genetic architecture between these populations, even though most observed PLPs are unique to a single cohort (96.4%, 54.2%, 52.8% of the PLPs in the UK, Dutch and Estonian cohorts, respectively) (**Supplementary data Fig. 16**). The proportion of PLPs in the UK Biobank cohort that was also found in the Dutch cohort is only 2.4% (1,338/54,758). Conversely, the proportion of PLPs in the Dutch cohort that was also present in the UK biobank cohort is 35.8% (1,338/3,734). This difference reflects the larger cohort size of the UK biobank cohort which is ∼100 times larger than the Dutch cohort. These results suggest that the similarity of the genetic landscapes as represented by the CR and AF is not due to the presence of shared PLPs, but more likely reflects selection acting independently in each of these populations.

## Discussion

We show that, at biobank scale, carriers of heterozygote PLPs in recessive constrained genes have reduced reproductive fitness, expressed as being more likely to be childless. By convention, the terms dominant and recessive are defined by the clinical consequences in heterozygotes, i.e. phenotypic expression in dominant conditions and no phenotype in recessive conditions^20^. This dichotomy is clearly simplistic. In dominant conditions such as achondroplasia, the biallelic state results in more severe phenotypes. Conversely, clinical effects have been observed in heterozygotes for various recessive disorders such as sickle cell trait, or Parkinson’s disease in carriers of Gaucher disease^21^. The phenotypic impacts of the genetic burden that we observe for heterozygous recessive variants align with proposals that the labels dominant and recessive disease should be seen as the extremes of a more graded scale of phenotypic expression^2^. We note that, while a recent analysis examined fitness effects for loss-of-function variants across the human genome, our findings are specific to PLPs in recessive genes, and the selective effect that we observe for recessive disease genes is not driven by the same LoF variants^13^.

Furthermore, we find that only carriers of PLPs in recessive ID genes have significantly fewer years of education. Whereas educational attainment is an obvious phenotype to assess in the context of ID genes, fitness of PLP heterozygotes in both ID and non-ID recessive genes may well be affected by other phenotypes. In fact, genes and variants that are associated with neurodevelopmental disorders, can also affect other organ systems^22,23^. For instance, heterozygosity for PLPs in *KDM5B*, a known recessive ID gene, has pleiotropic effects not restricted to cognitive function^22^. We indeed also observe an association of genetic burden in recessive ID genes and a higher number of ICD-10 diagnoses.

While our findings do not prove causality, the observed reproductive and phenotypic effects, associated with high genetic burden in recessive ID genes, are consistent with previous sociodemographic evidence for cognitive and behavioral effects as potent mediators of increased childlessness^16,19^. Although an increase in childlessness indicates selective effects on genetic variants, this does not mean that such rare recessive genetic variants carry any predictive value for an individual’s reproductive success. In fact, medical conditions, societal and secular trends, as well as the polygenic background are much more important^24^. Nonetheless, because genetic selection works over evolutionary time, small effects for the individual can add up over many generations and thereby modify the genetic landscape^8^.

Consistent with this, we find that the landscape of PLPs in recessive ID genes is relatively depleted for more common PLPs. This is in spite of the fact that approximately 1 in 5 (22%) of individuals in the UK Biobank is heterozygous for a PLP in a recessive ID gene, similar to what was found in smaller Dutch and Estonian cohorts^9^. Previous work supports the notion that recessive ID genes may be depleted for more common deleterious variants^3,25,26^. In line with this, studies in neurodevelopmental patient cohorts show only a small contribution of biallelic recessive pathogenic variants in non-consanguineous families^3,25,26^. Indeed, recessive inheritance remains the exception even for those non-consanguineous families where multiple siblings are affected with intellectual disability or severe autism^27,28^.

We show that while the genetic landscape per gene is very similar for recessive diseases, the allelic landscape is quite different for the three European populations (UK, Dutch and Estonian) studied here. This has practical implications, as international databases of pathogenic variants will often be of limited use if the pathogenic variants mostly remain local rather than global. Our results therefore underline the need for local and national databases of genetic variation in order to better understand and diagnose recessive disease in a population.

Since the genetic burden score represents the effect on heterozygotes, the selective effect for recessive genes should be less than for dominant genes. Indeed, the effect of LoF variants in non-recessive constrained genes on childlessness is detectable in ∼112,000 individuals, whereas for constrained recessive genes it was detectable from about ∼376,000 individuals onwards (**Supplementary Methods, Supplementary data Fig. 4**). It is therefore not surprising that we did not find sex-specific differences that were observed in previous studies. Similarly, we found that the associations with childlessness and cognitive phenotypes are only significant for the subset of ID genes (n=357). However, we expect that these associations may well be driven by a broader range of genes with modest effects, and that we currently lack the sample size to detect these.

In future studies, it will be important to assess additional phenotypes in heterozygotes, and to study non-European cohorts that include diseases such as hemoglobinopathies, which are underrepresented in our study. Furthermore, it is important to replicate these results in other European and non-European populations.

## Methods

To test whether there is selection against carriers of heterozygous pathogenic/likely-pathogenic variants (PLPs) in autosomal-recessive (AR) genes, we extracted genetic and phenotypic data from the UK Biobank^1^ (age, number of children, educational attainment, etc.; **Supplementary Table 14**) and used multiple generalized linear regressions to examine if there is an association between genetic burden, based on the PLPs harbored by an individual, and the tested phenotypes (**Supplementary Table 15**).

### Dataset

We accessed the UK Biobank’s final release of whole-exome sequencing data from July 2022 covering 469,787 participants via the Research Analysis Platform. Using GRCh38-aligned population-level variant call format files (data-field 23157) we extracted relevant variants and identified PLPs for further analysis.

We also downloaded the phenotypic data of interest (**Supplementary Table 14**) for all available participants. For this study, we only considered information gathered at the time of recruitment (which corresponds to instance 0 in the UK Biobank).

#### Sample selection

The initial cohort of 469,787 UK Biobank participants, excluding the participants who withdrew their consent (as of May 2023), was filtered for ancestry and relatedness, leaving 378,751 participants.

Ancestry was determined by the UK Biobank’s data-field 22006, which indicates samples who self-identified as ‘White British’ and have a very similar genetic ancestry based on a principal components analysis of their genotypes. Using this information 409,520 samples were selected for further analysis.

We have downloaded data on the relationships between UK Biobank participants by following the instructions outlined in resource 668. This dataset contains a pairwise list of individuals along with estimated kinship coefficients for each pair. Out of 409,520 European samples we excluded 30,769 samples consisting of one individual from each pair found to be second degree relatives or closer based on the kinship coefficient.

All subsequent analyses were based on these 378,751 unrelated samples of European ancestry.

To eliminate possible effects of compound heterozygous or homozygous variants in regression analyses, we additionally excluded 2,143 samples containing compound heterozygous or homozygous variants in 1,929 recessive genes.

#### Recessive genes selection

We selected a total of 1,929 genes (6,011 transcripts) with recessive phenotypes, as described by Fridman et al.^9^ (**Supplementary Table 1**). Briefly, OMIM genes associated with only recessive phenotype(s) were chosen for further analysis (1,605 genes). Genes linked with both autosomal-dominant (AD) and -recessive (AR) phenotypes were further assessed by their gnomAD^29^ pLI score. AD-AR genes were only included if their pLI score was lower than the 95th percentile of pLI score calculated for 930 manually curated AR-only genes causing severe phenotypes^30^ (pLI score ≤ 0.86), yielding 324 genes for inclusion^9^.

We additionally compared the distribution of s_het_ scores between 324 AD/AR and 1605 AR genes and confirmed that on average they have similar s_het_ scores (0.036 and 0.037 respectively; Welch’s t-test statistic: 0.53, p-value: 0.59).

#### PLP selection process from WES data

### Quality control

To ensure good sequencing and genotyping quality, we selected variants within the target region used by the WES capture experiment (UK Biobank resource 3803). Additionally, we only retained variants with at least 15x coverage (as specified by ‘DP’) for more than 90% of the cohort. This part of the variant selection process was performed on the DNAnexus JupyterLab with Spark cluster (v1.0.1) using Hail (v0.2.78). Thus we downloaded 1,544,703 variants located in the above-mentioned 1,929 recessive genes with at least one non-reference allele, together with associated genotypes.

### Evaluating variant outcomes

Further variant selection was performed as described by Fridman et al.^9^ with some modifications (**Supplementary Fig. 1**). We used the list of transcripts from the HGMD (http://www.hgmd.cf.ac.uk/ac/index.php) database (v2018.3) as a reference. The variant outcomes were extracted using VEP^31^(v104). If a gene had only one transcript described in the HGMD database, the outcome of the variant was defined based on this transcript. For genes with several transcripts, agreement of >50% on the variant outcome (LoF/missense/other) was necessary for the outcome to be chosen. For all other cases (24,998 variants) the most severe outcome was considered, of which 2,032 (8.1%) passed the selection process and were included in the PLPs list.

### Indels filtering

In order to prevent a high incidence of false-positive indels, we adjusted the selection process by using indels in AD genes associated with intellectual disability (AD-ID) as a proxy for false positives, as described in Fridman et al.^9^ (**Supplementary Fig. 1**).

### Pathogenic and likely pathogenic variants (PLPs) selection process

We created a list of presumable PLPs in our cohort of 378,751 individuals. First, we filtered out variants with ≥5% heterozygote or ≥1% homozygote frequency within the cohort. We then selected only variants that met at least one of three criteria (**Supplementary Fig. 1**):

1. Classified as PLP by ClinVar (https://www.ncbi.nlm.nih.gov/clinvar/) (downloaded 30.4.2023) with a review status of ≥2 stars, or classified as PLP by the VKGL^25^ database (v. April 2021). This curated database is publicly available and comprises DNA variant classifications established based on (former) diagnostic reports of all 9 Dutch accredited laboratories;
2. Rare loss-of-function (LoF) variants (nonsense, frameshift, canonical splicing) with <1% or unknown frequency in gnomAD^29^. Additional filtering was performed for indels as described in **Supplementary Fig. 1**;
3. Missense variants classified as PLP by ≥2/3 databases: InterVar^32^ (an automated ACMG classifier, downloaded 30.11.2022), ClinVar with a review status of <2 stars, and HGMD (http://www.hgmd.cf.ac.uk/ac/index.php) (indicated as a disease-causing variant by the DM flag).

Variants from criteria 2 and 3 were excluded if they contradicted the first criterion, i.e. were classified as benign/likely-benign by Clinvar with a review status of ≥2 stars or by the VKGL^33^. For the AD-AR genes, only LoF variants were included in the final PLPs list.

In total, the selection process filtered out ∼95% of the initial number of variants in the selected regions.

Additionally, all of the frequency drop-outs, i.e. variants with ≥5% heterozygotes or ≥1% homozygotes frequency within the cohort, were filtered by the same selection process, and the one variant that passed the selection process was manually curated. The manual curation led to the re-inclusion of this variant with >5% carrier frequency, *HFE* [MIM 613609] p.Cys282Tyr, for which there are 55,998 (13.8%) heterozygotes and 2,524 (0.6%) homozygotes in the cohort.

Overall 54,758 variants were classified as PLPs, including 30,405 substitutions (55.5%), 16,487 deletions (30.1%), and 7,866 insertions (14.4%). From this list, 142 variants in 115 genes were seen in a homozygous state in a range of 1-2524 individuals (average 21 individuals, median 1). We further inspected the PLP counts for each sample and found no outliers (**Supplementary Table 16**).

For the regression analyses we considered a smaller set of 50,568 rare PLPs that were observed no more than 20 times in homozygous state and never in heterozygous state.

### Validation and manual curation

Manual curation was performed for PLPs with heterozygote frequency ≥0.5% and/or with five or more homozygotes in the cohort. Out of 54,758 PLPs found in the cohort, 34 (0.06%) PLPs met the frequency criteria for manual curation. It showed that 32/34 variants were previously described, and that the other 2 variants are rare LoF variants not reported previously on ClinVar. Furthermore, 13/34 variants have been reported to cause only a mild phenotype or even appear asymptomatic when seen in a homozygous state. For the CR calculations, we only considered couples who can potentially have a compound heterozygous child for a severe and mild variants.

#### LoF singleton carriers selection

To eliminate the possibility that the observed effect of PLPs in recessive genes could be confounded by pathogenic variants in other genes, an additional dataset with carriers of singleton LoF variants was collected from the UK Biobank. For the purposes of this analysis, we only considered LoF variants located in 1,983 autosomal dominant and recessive genes under stronger selection, as specified by s_het_ ^15^ ≥ 0.15.

The selection procedure for the variants was similar to Gardner et al^13^. Genotypes were set to “null” if DP<7 or GQ<20 or the binomial test P-value ≤ 0.001 for alternate versus reference reads on heterozygous genotypes only. We filtered out variants if more than 50% of genotypes were missing.

All variants were annotated with VEP^31^ version 104. Variants were considered LoF if they were annotated as having splice donor/acceptor, stop gained, or frameshift consequence in the canonical transcript. We only considered high-confidence variants as annotated by LOFTEE^29^. This yielded 25,441 unique LoF variants, out of which 16,975 were singleton (occurred only once in the cohort). 1,024 were located in 1,929 recessive genes and 15,951 in the rest.

#### Synonymous variants selection

As most of the synonymous variants are neutral or only slightly deleterious, they are unlikely to have phenotypic or selective effects, making them suitable as negative controls.

We selected all rare (allele count (AC) ≤ 20) synonymous variants (n=247,665) in a set of 1,929 recessive genes that were not observed in a homozygous state, thus replicating the allele frequency criteria for PLPs. We sampled from this set of synonymous variants 300 times to match the number of variants observed for PLPs (n=50,568).

For each variant set we performed generalized linear regressions as described below for the phenotypes of interest. This approach allowed us to generate sampling distributions of the estimated effects of synonymous variants on the selected phenotypes.

#### Phenotype selection

Phenotypic data was chosen based on our interest in fitness effects and education as a proxy for cognitive ability.

The list of all the phenotypes and the processing made for each phenotype are described more in detail in **Supplementary Table 14**.

1. Fields “Number of children fathered” (field number 2405) for males and “Number of live births” (field number 2734) for females were used to determine the number of children of each participant and also the “childlessness” status in case the number was 0. We did not account for cases of miscarriages or stillbirths in females due to the lack of this information for their male partners.
2. The “Qualification” field (field number 6138) was used to determine educational attainment. The highest qualification reported was mapped to the International Standard Classification for Education (ISCED) coding for years of education as described previously in the literature^34^.
3. The “Hair color” field (field number 1747) was chosen as a control phenotype as a phenotype that is not expected to be related to fertility or education. However, we acknowledge that no phenotype may be fully neutral with respect to education and reproductive success. There is evidence for an effect of the major hair and skin color MC1R locus on childlessness^35^ even though that effect appears not be mediated by the hair color phenotype.

#### Regressions

For the regression analysis, we focused on PLPs that were never observed in a homozygous state and occurred in a heterozygous state no more than 20 times within the cohort (0.005%). Based on the observed PLPs, we calculated individual genetic burden for the UK Biobank participants as described in Gardner et al.^13^

Briefly, we combined *s*_*het*_ scores of the genes in which an individual *i* has PLPs using formula: *s*_*het*_ [*i*] = 1 − ∏_*g*∈*𝒢*_ (1 − *s*_*het*_ [*g*]), where *g* denotes a gene and product iterates over all genes with PLPs from a gene set *G*.

We restricted variants to different sets of genes, such as the full AR 1,929 gene set or disorder group genes (e.g ID, Blindness, AR without ID, etc.), therefore calculating several genetic burden (*s*_*het*_ [*i*]) scores for an individual. Additionally, we used different sources for scores^15,36^ and repeated the analysis with pLI scores^29^ (**Supplementary Tables 3-6**).

Participants who carried PLPs in genes with no known scores were excluded from the corresponding analysis. We considered the genetic burden to be 0 if a participant did not harbor any PLPs in the selected genes.

We tested the association of genetic burden with phenotypes as proposed in Gardner et al.^13^ using the regression of the form:

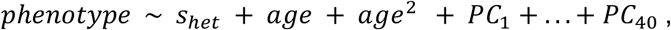

where *phenotype* and individual genetic burden *s*_*het*_ are determined by the context of the analysis, *PC*_1_, … , *PC*_40_ denote 40 genetic principal components. By using this form of regression we corrected for possible biases due to age and ancestry. We used binomial family distribution for binary phenotypes and Gaussian for continuous phenotypes.

When testing phenotype associations with the number of genes affected by PLP or carriership status, we use the same regression form replacing genetic burden by the corresponding explanatory variable.

All regressions were performed using generalized linear models from statsmodels 0.13.5 package in Python 3.7.6. For more information about the types of regressions and Bonferroni correction applied see **Supplementary Table 15**.

#### Testing the equivalence of the association strength

In order to identify whether the association strength of different variants or sets of genes is the same (for example, PLPs in recessive genes vs singleton LoFs in non-recessive highly constrained genes or PLPs in recessive ID genes vs PLPs in other recessive genes), we performed the regression on this phenotype adding both genetic burdens as predictors. Since genetic burdens were calculated using different gene sets, their influence on the phenotype of interest is independent. Therefore, the regression coefficients obtained by this approach are similar to what would be obtained by running regressions with two genetic burdens separately. We used a Wald test for constraints to check the linear hypothesis, that both regression coefficients associated with genetic burdens are equal.

To compare the differences in association strength with phenotypes between females and males we incorporated an interaction term between the genetic burden and sex. We conclude that there is a significant difference between males and females if the interaction term is significantly different from zero as reported by the regression term’s p-value.

#### Simulated recessive ID and non-ID gene sets with matching s_het_ distributions

To separate the effects of recessive ID genes from those associated with high s_het_ enrichment, we compared the ID gene set with a gene set of similar s_het_ distribution, excluding ID genes. We created bins for s_het_ per gene in 0.05 increments and sampled a number of non-ID genes per bin equal to the number of ID genes. This process generated new gene sets with matching gene counts and s_het_ distributions as the ID panel. We repeated this procedure 20 times, resulting in 20 distinct recessive non-ID gene sets.

We run generalized linear regressions to compare the association strength with the phenotypes of interest between these 20 artificial gene sets and recessive ID genes.

#### Simulations of the reduced cohort size

To estimate the required cohort size for detecting observable effects, we conducted simulations by reducing the cohort size in increments of 10%, ranging from 10% to 90% of the original dataset. For each cohort size, individuals were randomly sampled 20 times without replacement and the analysis was performed for the association of genetic burden with childlessness for PLPs in recessive genes and singleton LoF in non-recessive highly constrained genes. This approach yielded distributions of odds ratios and p-values for each reduced cohort size and variant set.

#### Gene panels and consanguinity ratio (CR)

Genes were divided into non-overlapping disorder groups according to their related phenotypes, as described in Fridman et al.^9^ (**Supplementary Table 1**).

CR was calculated for the 1,929 AR gene set and for each disorder group, as the fold-increase in at-risk couples (ARCs) due to first-cousins consanguinity when compared to non-consanguineous couples.

We first verified that the population adheres to the Hardy-Weinberg equilibrium. We compared the observed number of heterozygotes (Aa) to the expected number (2pq) using a chi-squared test, showing that variants adhere to the Hardy-Weinberg equilibrium and there is no significant difference between observed and expected with a P-value of 0.4.

Next, we defined *p*_s_ and *p*_*m*_ as the frequencies of severe and mild alleles (see “Validation and manual curation” section) respectively in a particular gene. Following the Hardy-Weinberg equilibrium frequencies, we could establish frequencies for heterozygotes as 2*p*_s_ *q*_s_ and 2*p*_*m*_ *q*_*m*_. Using these frequencies, the number of ARCs for consanguineous (first cousins) and non-consanguineous cases per gene could be calculated as following, accounting for ARCs for both homozygous and compound heterozygous state for severe alleles, and compound heterozygous state of severe-mild alleles:

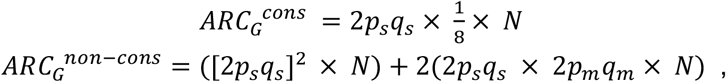

where *N* is a number of individuals in the cohort and denotes a particular gene. Therefore, the ratio between ARCs for consanguineous versus non-consanguineous cases for a set of genes could be calculated as:

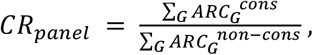

where summation iterates over all genes in the panel.

To examine whether the differences between the CR scores among the disorder groups are statistically significant, we ran 10,000 simulations per disorder group. In each simulation, we randomly assigned genes into panels of the same total coding length as the original panels. We computed the p-values based on how many times the simulated CR was different from the actual CR for each disorder group using a permutation test with a Bonferroni correction for 13 tests (**Supplementary Table 12**).

## Data availability

The raw data used in this study are available as part of the UK Biobank dataset.

## Code availability

The code used to generate the data for this project is available on GitHub: https://github.com/Genome-Bioinformatics-RadboudUMC/ukbb_recessive_public

## Acknowledgments

We thank Eugene Gardner, Juliet Hampstead , Hilary Martin and Matthew Hurles for fruitful discussions; Shai Carmi for advising about population genetics matters. This research has been conducted using the UK Biobank Resource under application number 66493. This project was financially supported by a VIDI grant from the Dutch Research Council (917-17-353 to C.G.) and an AI for Health PhD grant from Radboudumc. This project was supported by a gift from the Koum Foundation (to E.L.L.). E.L.L. is Robin Chemers Neustein Director of Medical Genetics.

## Author contributions

H.G.B, C.G. and E.L.L. supervised the study, H.F. and G.K. developed a data collection pipeline, statistical methods and analyzed data. H.F., G.K., H.G.B, C.G and E.L.L. wrote the manuscript.

