## Supplementary material for "Reproductive and cognitive effects in carriers of recessive pathogenic variants": Supplemenary Figures

### Supplementary data figures

**Supplementary data Fig. 1.** Variant and carrier frequency of heterozygous PLPs.

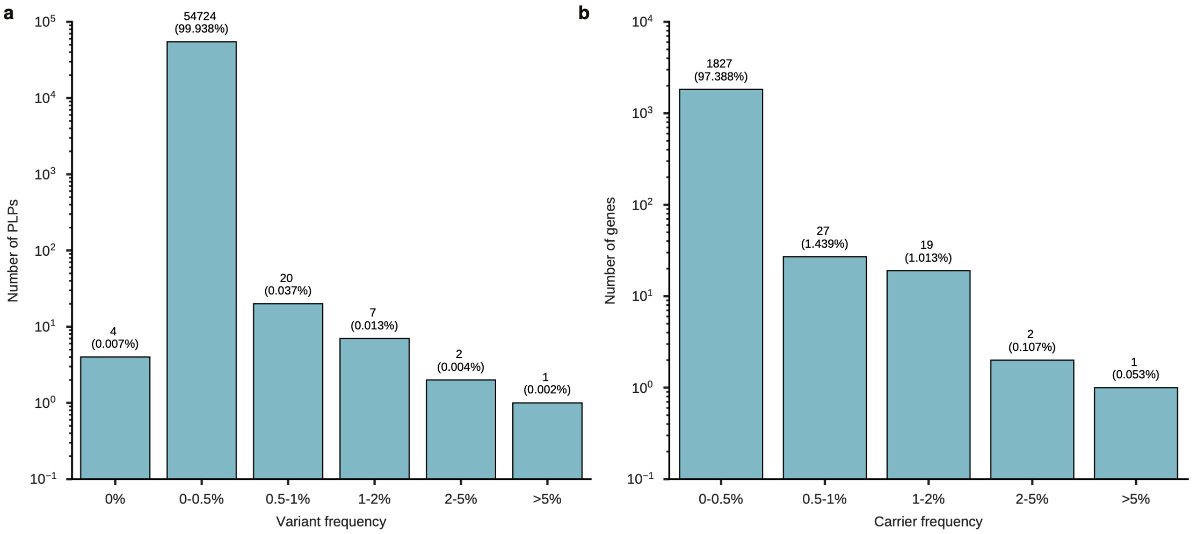


**(a)** Distribution of carrier frequency of heterozygous PLPs. Four variants were never observed in the heterozygous state, but only once in the homozygous state.

**(b)** Distribution of per gene carrier frequency of heterozygous PLPs.

**Supplementary data Fig. 2.** Association of genetic burden for recessive disease with childlessness for different constraint scores.

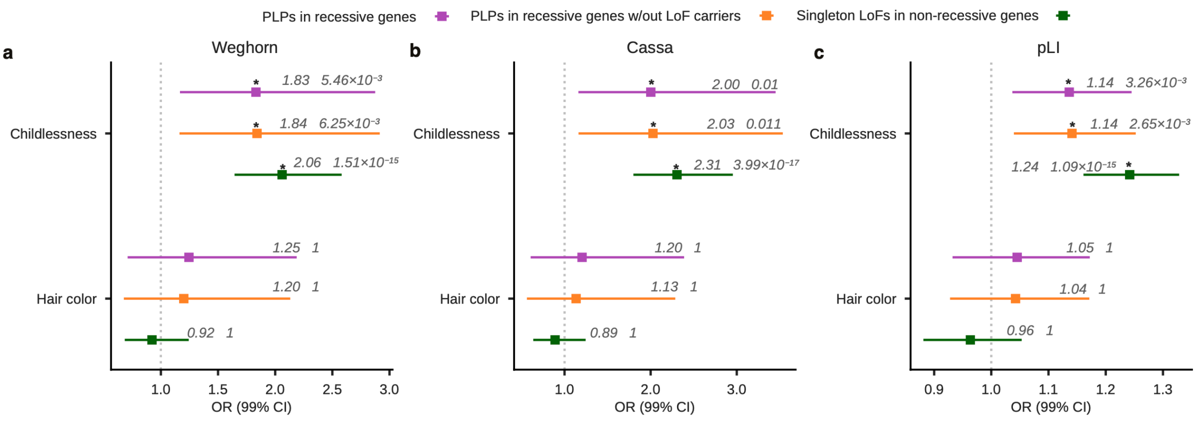


Effect of genetic burden on childlessness and hair color (as a control phenotype) for heterozygous PLPs in recessive genes (purple), PLPs in recessive genes excluding carriers of LoF in non-recessive genes (orange), and singleton LoFs in highly constrained (s-het > 0.15) non-recessive genes (green). Colored lines indicate odds ratios for the phenotypes with 99% confidence intervals; dashed gray line indicates an odds ratio of 1; p-values are adjusted for multiple testing using Bonferroni correction; significant associations are marked with an asterisk. The results are for three different s-het scores: Weghorn **(a)**, Cassa **(b)** and pLI **(c)**.

**Supplementary data Fig. 3.** Distribution of the number of rare PLPs and genetic burden per individual stratified by childlessness.

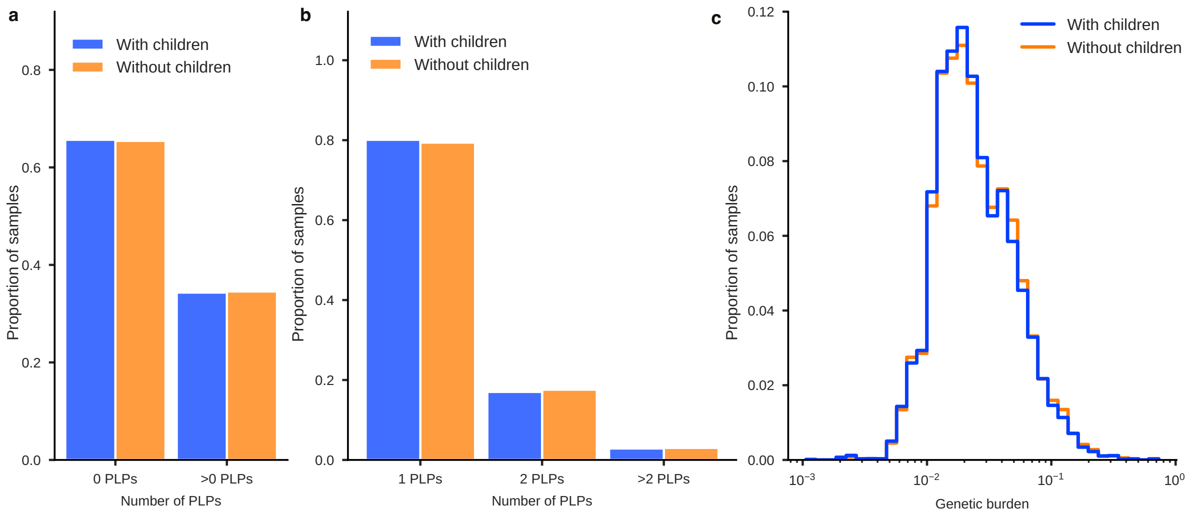


**(a)** Proportion of individuals with/without any PLPs for individuals with (blue) and without (orange) children.
**(b)** Distribution of the number of PLPs per individual with (blue) and without (orange) children.
**(c)** Distribution of the genetic burden per individual for individuals with (blue) and without (orange) children.

**Supplementary data Fig. 4.** Simulations of the effect of sample size on detection of the association with childlessness.

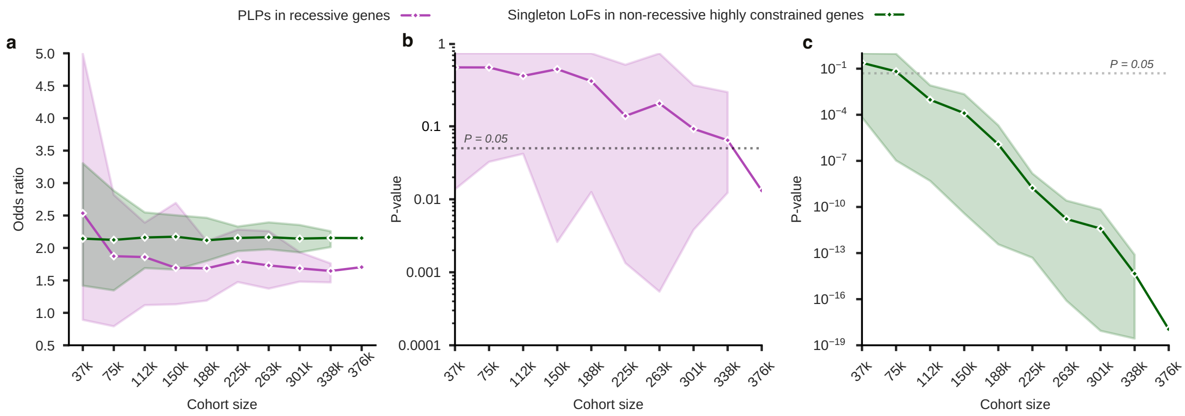


Simulated estimation of the odds-ratios **(a)** and corresponding p-values (**b**)-(**c**) depicted with a spread (95% percentile interval for n=20 simulations per cohort size) for the effect of cohort size on the association of childlessness with the genetic burden of PLPs. Shown are simulation results for heterozygous carriers of PLPs in recessive genes (purple) and of carriers of singleton LoFs carriers in non-recessive highly constrained genes (green). The dotted gray line marks the significance level of P=0.05.

**Supplementary data Fig. 5.** Association of genetic burden for recessive disease with educational attainment, fluid intelligence score, and log-transformed number of ICD-10 diagnoses for different constraint scores.

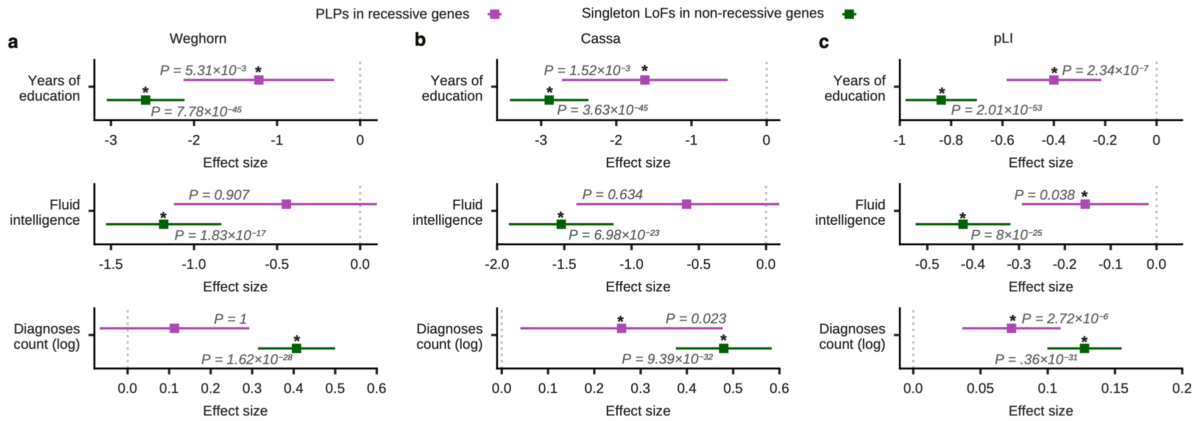


Effect of genetic burden on educational attainment (measured in years of education), fluid intelligence score and log-transformed number of ICD-10 diagnoses for PLPs in all recessive (purple) genes and singleton LoFs in non-recessive highly constrained (green) genes. Colored lines indicate effect sizes (estimated regression coefficients) with 99% confidence intervals; dashed gray line indicates the effect size of 0; p-values are adjusted for multiple testing using Bonferroni correction; significant associations are marked with an asterisk. The results are for three different s-het scores: Weghorn **(a)**, Cassa **(b)** and pLI **(c)**.

**Supplementary data Fig. 6.** Genetic burden and childlessness: controlling for the number of ICD-10 diagnoses, infertility, and having a partner.

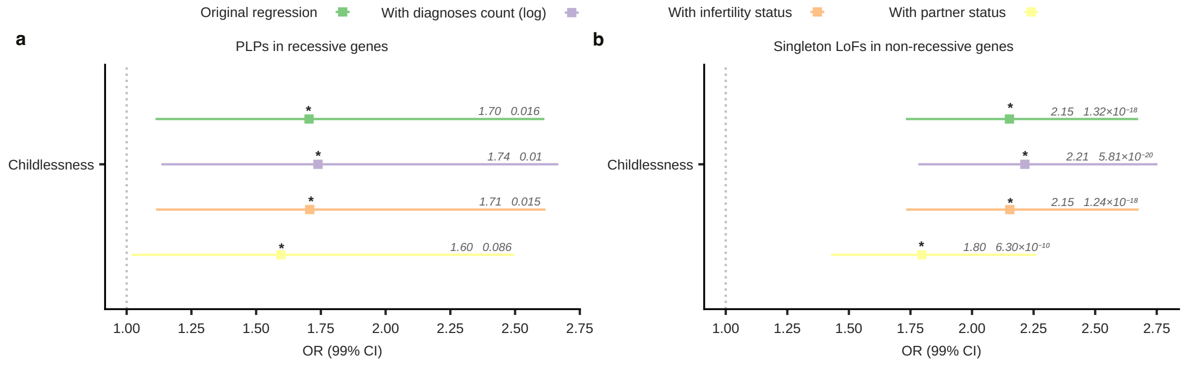


Effect of genetic burden on childlessness for heterozygous PLPs in recessive genes. Shown are results of the original analysis (green), and of controlling for the number of ICD-10 diagnoses (purple), infertility (orange), or having a partner (yellow). Results are shown for PLPs in recessive genes (**a**) and singleton LoFs in non-recessive highly constrained genes (**b**). Colored lines indicate odds ratios for childlessness with 99% confidence intervals; dashed gray line indicates odds ratio of 1; p-values are adjusted for multiple testing using Bonferroni correction; significant p-values are marked with an asterisk.

**Supplementary data Fig. 7.** Genetic burden associations: controlling for deprivations.

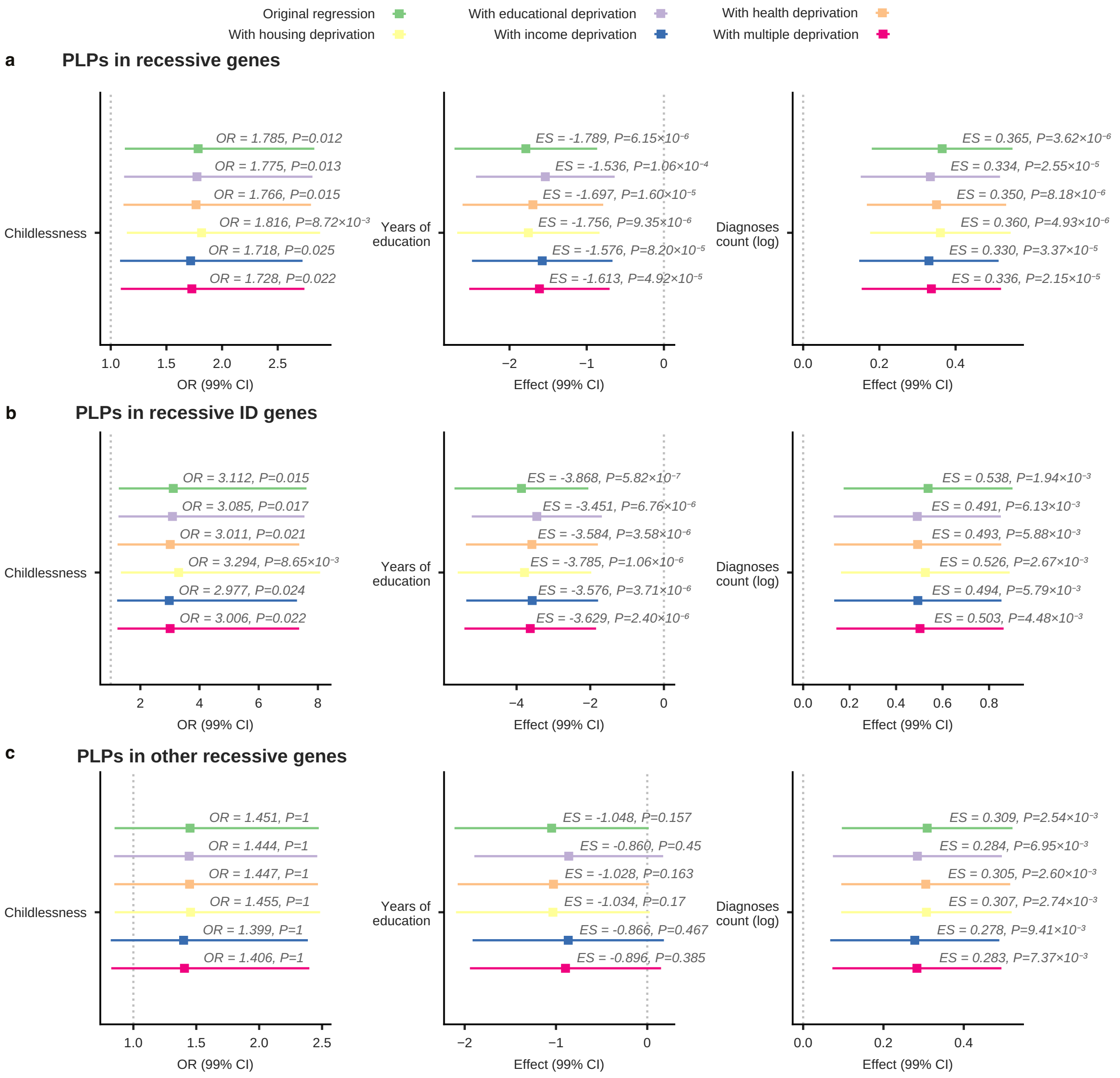


Effect of genetic burden on childlessness, educational attainment (in years of education), and log-transformed number of ICD-10 diagnoses for heterozygous PLPs in all recessive genes (**a**), recessive ID genes (**b**) and all other recessive genes (**c**). The original regression is shown in green, controlling for educational deprivation in purple, health deprivation in orange, housing deprivation in yellow, income deprivation in blue, and multiple deprivations in magenta (Supplementary table 14). Colored lines indicate odds ratios for childlessness or effect sizes (estimated regression coefficients) with 99% confidence intervals; dashed gray line indicates an odds ratio of 1 (for childlessness) or effect size of 0; p p-values are adjusted for multiple testing using Bonferroni correction.

**Supplementary data Fig. 8.** Association of genetic burden with various phenotypes for PLPs in recessive ID genes and all other recessive genes, using different constraint scores.

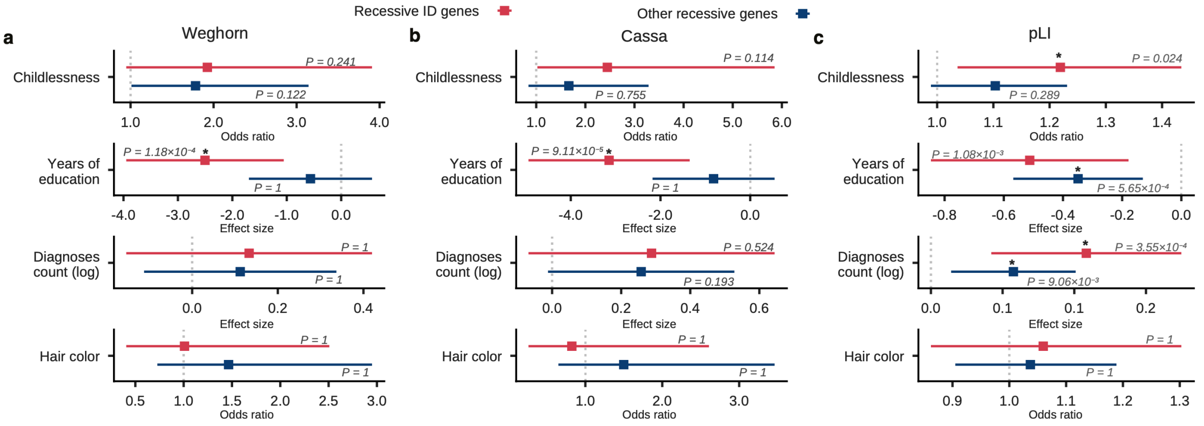


The comparison between PLPs in recessive ID genes (red) and all other recessive genes (blue) for the effects of genetic burden on childlessness, educational attainment (measured in years of education), log-transformed number of ICD-10 diagnoses, and hair color (as a control phenotype). The results are for three different s-het scores: Weghorn **(a)**, Cassa **(b),** and pLI **(c)**. Colored lines indicate odds ratio (for childlessness and hair color) or effect sizes (estimated regression coefficients) with 99% confidence intervals; dashed gray line indicates the odds ratio of 1 (for childlessness and hair color) or the effect size of 0; p-values are adjusted for multiple testing using Bonferroni correction; significant associations are marked with an asterisk.

**Supplementary data Fig. 9.** Comparison of the effects between synonymous variants and PLPs in recessive ID genes and all the other recessive genes.

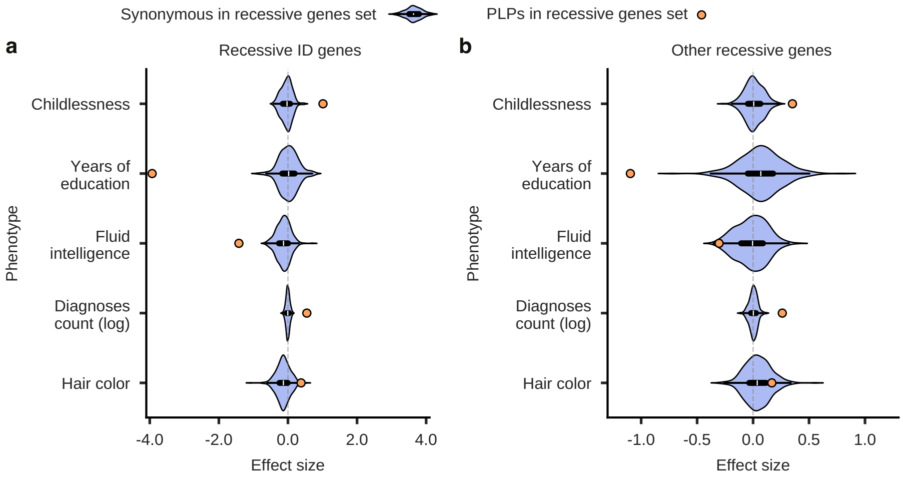


Distribution of the estimated effect sizes (regression coefficients) on the studied phenotypes for synonymous variants in recessive genes re-sampled 300 times (**Methods**; light purple violin plots) compared with the estimated effect size for PLPs (orange dots) in (**a**) recessive genes for ID and (**b**) all other recessive genes.

**Supplementary data Fig. 10.** Roulette $s_{het}$ distribution over different gene sets.

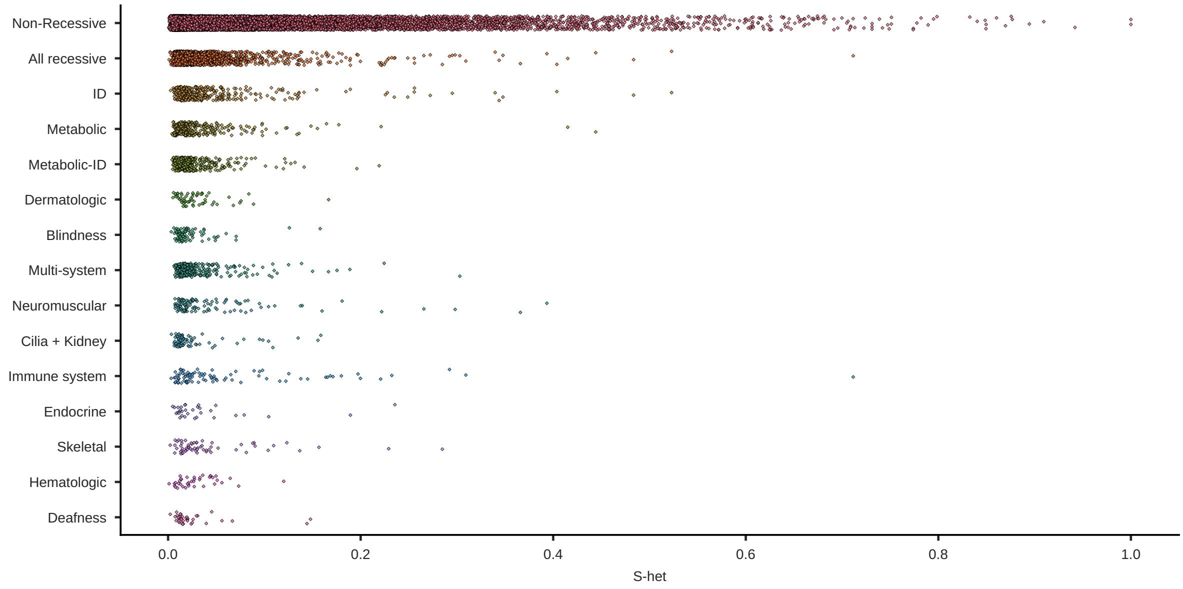


Roulette $s_{het}$ distribution in different gene sets. Each dot represents a particular gene. Genes are grouped by disease phenotypes (y-axis) with $s_{het}$ values on the x-axis.

**Supplementary data Fig. 11.** Sampling non-ID genes with the same s-het distribution as ID.

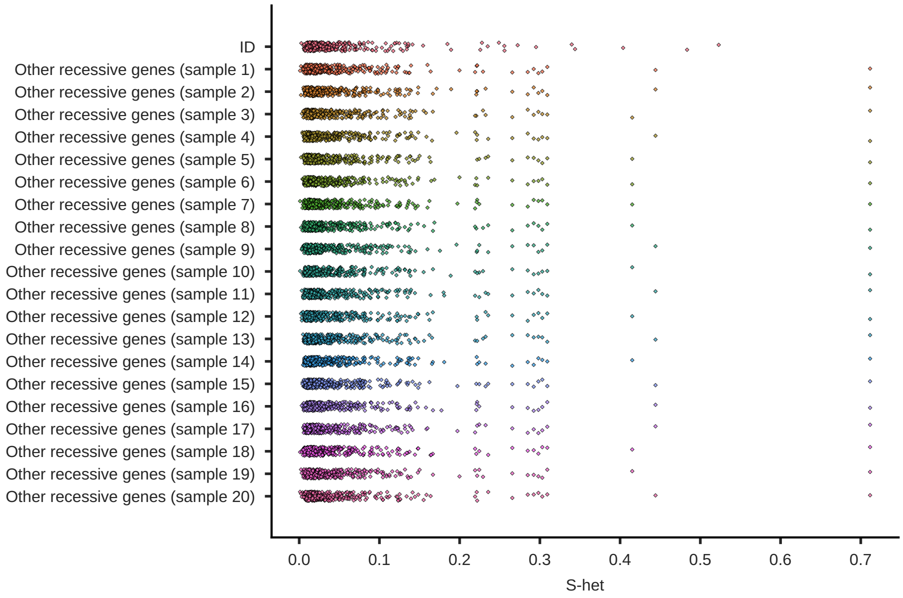


Roulette $s_{het}$ distribution over simulated gene sets with a distribution matching ID set (**Methods**). Each dot represents a particular gene. Simulated gene sets are shown on the y-axis. Values are scattered across the x-axis.

**Supplementary data Fig. 12.** Genetic burden associations in a sampling of the non-ID genes with the same $s_{het}$ distribution as ID genes.

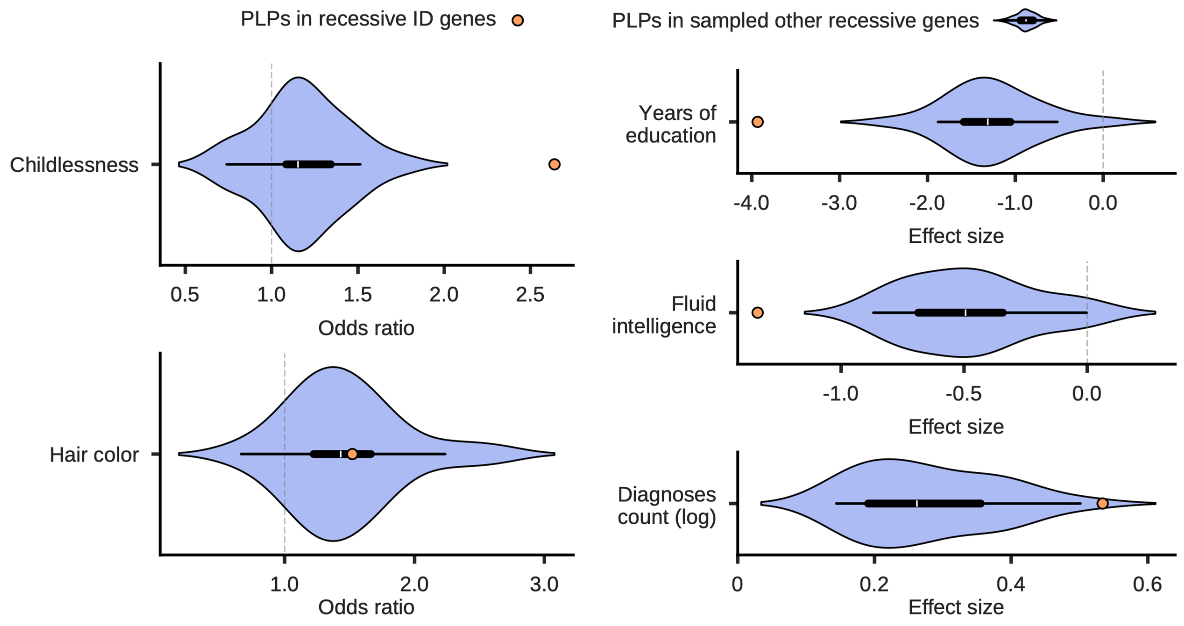


Distribution of the estimated odds ratios (for childlessness and hair color) and effect sizes (regression coefficients) for PLPs in simulated non-ID recessive gene sets (**Methods**; light purple violin plots) re-sampled 20 times to match the s-het distribution of the recessive ID gene set (orange dots).

**Supplementary data Fig. 13.** Sex-specific effects.

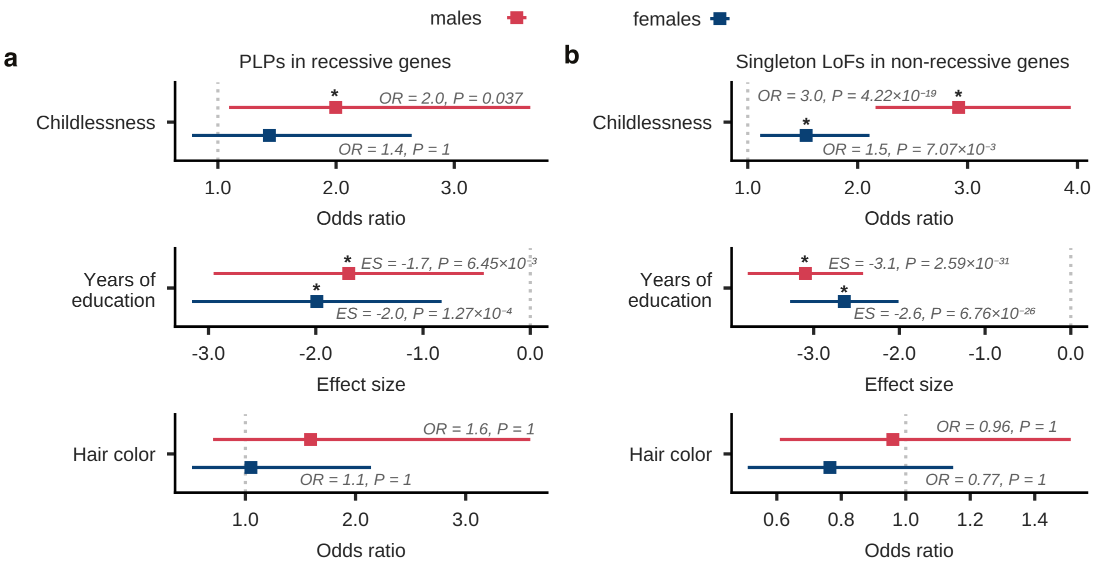


Effects for males (red) and females (blue) of genetic burden on various phenotypes for heterozygous PLPs in recessive genes (**a**) and singleton LoFs in non-recessive highly constrained genes (**b**). Colored lines indicate odds ratio (for childlessness and hair color) or effect sizes (estimated regression coefficients) with 99% confidence intervals; dashed gray line indicates the odds ratio of 1 (for childlessness and hair color) or the effect size of 0; p-values are adjusted for multiple testing using Bonferroni correction; significant associations are marked with an asterisk.

**Supplementary data Fig. 14.** Consanguinity ratio scores (CR) for different disorder groups in three European populations.

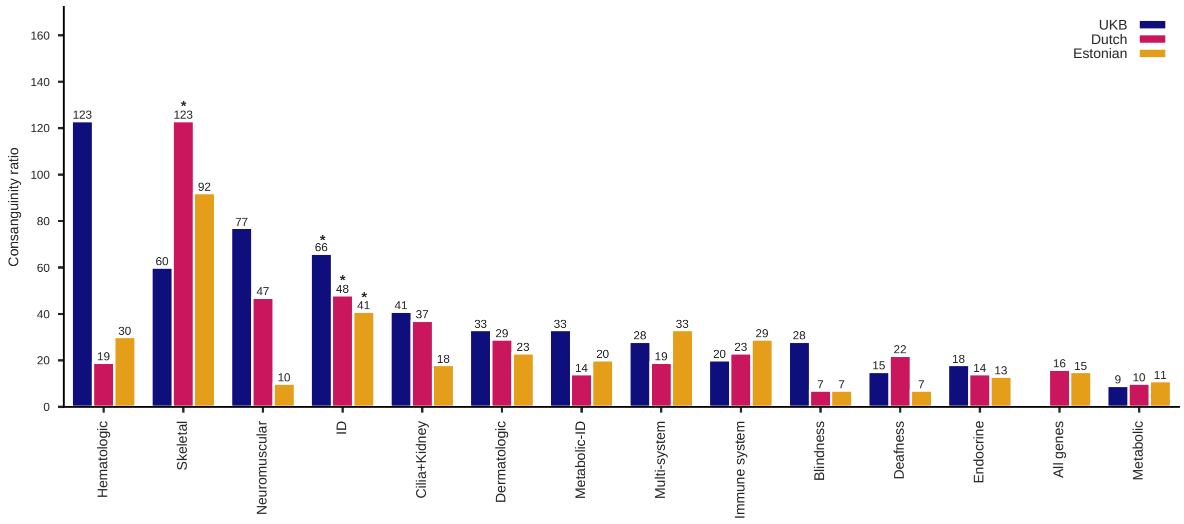


Consanguinity ratio scores (CRs) for 13 disorder groups calculated for 3 European populations: UK Biobank (blue), Dutch (magenta) and Estonian (orange) cohorts. Scores for the Dutch and Estonian cohorts are taken from Fridman et al.[9]; Scores marked in asterisk are significantly higher than seen in a random set of recessive genes with the same coding length (Methods). We note that CR scores for some of the other disorder groups (Skeletal, Neuromuscular, Hematologic) are similar to or higher than the CR score obtained for ID genes, yet these did not reach significance.

**Supplementary data Fig. 15.** Correlation of allele frequencies in disease categories between a Dutch cohort and the UK Biobank.

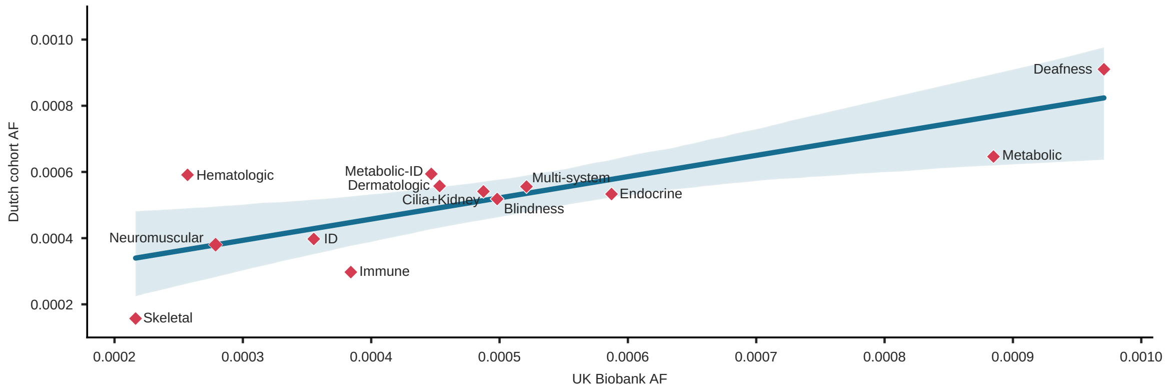


Correlation of average PLP allele frequencies (AF) per disorder group between the UK Biobank and Dutch cohorts with 95% confidence interval for the regression estimate (derived from n=1000 bootstrap samples). The X-axis shows the AF in the UK Biobank cohort, Y-axis shows the AF in the Dutch cohort. Pearson correlation coefficient: 0.79, p-value: 0.001. AF for the Dutch cohort are taken from Fridman et al.[9]

**Supplementary data Fig. 16.** PLPs overlap in three European cohorts.

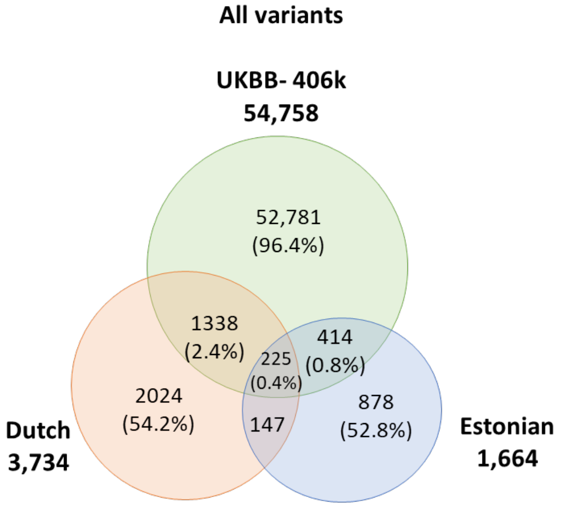


Overlaps of the number of PLPs in UKB (green), Dutch (orange), and Estonian (blue) cohorts, were calculated for all PLPs. Data on the Dutch and Estonian cohorts are taken from Fridman et al.[9]
