## Supplementary material for "Reproductive and cognitive effects in carriers of recessive pathogenic variants": Supplemenary Tables

### Supplementary Tables

**Supplementary Table 1-** analyzed genes (Excel)

**Supplementary Table 2.** Regression performed on all recessive genes using several sources for genetic burden scores.

Genetic burden scores:

- Roulette-based scores from Seplyarskiy et al.
- Weghorn et al. using a demographic model that includes genetic drift
- Cassa et al.
- pLI scores from gnomAD v.2.1.1.

* Significant P-values are marked in bold and with an asterisk

| Genetic burden | Phenotype | Odds ratio (OR)/ effect size (ES) | P-value | Corrected p-value | Observations |
|---|---|---|---|---|---|
| Roulette | **childlessness** | OR = 1.7 | 0.00132 | **0.0132*** | 368,224 |
| | **years of education** | ES = -1.856 | $2.65\times10^{-8}$ | **$2.65\times10^{-7}$*** | 366,520 |
| | **log(# of ICD-10 diagnoses)** | ES = 0.33 | $5.73\times10^{-7}$ | **$5.73\times10^{-6}$*** | 369,994 |
| | infertility | OR = 0.51 | 0.638 | 1 | 369,994 |
| | living with a partner | OR = 0.75 | 0.062 | 0.62 | 368,038 |
| | fluid intelligence score | ES = -0.533 | 0.029 | 0.29 | 118,556 |
| | hair color | OR = 1.24 | 0.297 | 1 | 369,270 |
| Weghorn | **childlessness** | OR = 1.832 | $5.46\times10^{-4}$ | **$5.46\times10^{-3}$*** | 353,425 |
| | **years of education** | ES = -1.220 | $5.31\times10^{-4}$ | **$5.31\times10^{-3}$*** | 351,777 |
| | log(# of ICD-10 diagnoses) | ES = 0.113 | 0.106 | 1 | 355,116 |
| | infertility | OR = 0.157 | 0.267 | 1 | 355,116 |
| | living with a partner | OR = 0.721 | 0.042 | 0.418 | 353,231 |
| | fluid intelligence score | ES = -0.444 | 0.091 | 0.907 | 113,714 |
| | hair color | OR = 1.246 | 0.316 | 1 | 354,424 |
| Cassa | **childlessness** | OR = 2.002 | $1.03\times10^{-3}$ | **0.01*** | 353,246 |
| | **years of education** | ES = -1.620 | $1.52\times10^{-4}$ | **$1.52\times10^{-3}$*** | 351,601 |
| | **log(# of ICD-10 diagnoses)** | ES = 0.259 | $2.28\times10^{-3}$ | **0.023*** | 354,935 |
| | infertility | OR = 0.496 | 0.707 | 1 | 354,935 |
| | living with a partner | OR = 0.683 | 0.05 | 0.497 | 353,057 |
| | fluid intelligence score | ES = -0.591 | 0.063 | 0.634 | 113,648 |
| | hair color | OR = 1.204 | 0.486 | 1 | 354,245 |
| pLI | **childlessness** | OR = 1.136 | $3.26\times10^{-4}$ | **$3.26\times10^{-3}$*** | 337,479 |
| | **years of education** | ES = -0.400 | $2.34\times10^{-8}$ | **$2.34\times10^{-7}$*** | 335,892 |
| | **log(# of ICD-10 diagnoses)** | ES = 0.073 | $2.72\times10^{-7}$ | **$2.72\times10^{-6}$*** | 339,097 |
| | infertility | OR = 0.737 | 0.35 | 1 | 339,097 |
| | living with a partner | OR = 0.933 | 0.032 | 0.32 | 337,301 |
| | **fluid intelligence score** | ES = -0.156 | $3.77\times10^{-3}$ | **0.038*** | 108,569 |
| | hair color | OR = 1.045 | 0.32 | 1 | 338,423 |

**Supplementary Table 3.**

Regression analysis for childlessness on:

- PLPs carriership status/number of genes affected by PLP for all 1929 recessive genes
- singleton LoFs carriership status for all non-recessive highly constrained ($s_{het}$ >0.15) genes

* Significant P-values are marked in bold and with an asterisk

| Variants | Phenotype | Odds ratio | P-value | Observations | Feature |
|---|---|---|---|---|---|
| PLPs in 1,929 AR genes | childlessness | 1 | 0.988 | 374,813 | heterozygosity status |
| | childlessness | 1 | 0.58 | 374,813 | number of genes with PLPs |
| singleton LoFs in non-AR highly constrained genes | **childlessness** | 1.19 | **$5.41 \times 10^{-14}$*** | 374,813 | heterozygosity status |

**Supplementary Table 4.** Regressions performed on all recessive genes using different sources for genetic burden scores excluding heterozygotes for singleton LoF variants in non-AR highly constrained genes or adding heterozygosity as a covariate.
Genetic burden scores:

- Roulette-based scores from Seplyarskiy et al.
- Weghorn et al. using a demographic model that includes genetic drift
- Cassa et al.
- pLI scores from gnomAD v.2.1.1.

* Significant P-values are marked in bold and with an asterisk

| Genetic burden | Phenotype | Adding singleton LoF heterozygosity as a covariate | | | | Excluding singleton LoF heterozygotes | | | |
|---|---|---|---|---|---|---|---|---|---|
| | | Odds ratio | P-value | Corrected p-value | Observations | Odds ratio | P-value | Corrected p-value | Observations |
| Roulette | childlessness | 1.71 | 0.0013 | **0.013*** | 368,085 | 1.72 | 0.0013 | **0.013*** | 356,962 |
| | hair color | 1.25 | 0.276 | 1 | 369,131 | 1.21 | 0.371 | 1 | 357,972 |
| Weghorn | childlessness | 1.82 | 0.0006 | **$5.97\times10^{-3}$*** | 353,324 | 1.84 | 0.0006 | **$6.25\times10^{-3}$*** | 342,645 |
| | hair color | 1.25 | 0.3 | 1 | 354,323 | 1.2 | 0.4 | 1 | 343,611 |
| Cassa | childlessness | 1.99 | 0.0011 | **0.011*** | 353,123 | 1.99 | 0.0011 | **0.011*** | 342,495 |
| | hair color | 1.22 | 0.46 | 1 | 354,122 | 1.22 | 0.46 | 1 | 343,461 |
| pLI | childlessness | 1.14 | 0.00033 | **$3.30\times10^{-3}$*** | 337,433 | 1.14 | 0.000265 | **$2.65\times10^{-3}$*** | 327,226 |
| | hair color | 1.05 | 0.31 | 1 | 338,377 | 1.04 | 0.36 | 1 | 328,140 |

**Supplementary Table 5.** Regressions performed on singleton LoF variants in non-recessive highly constrained genes ($s_{het} \geq 0.15$) using several sources for genetic burden scores.
Genetic burden scores:

- Roulette-based scores from Seplyarskiy et al.
- Weghorn et al. using a demographic model that includes genetic drift
- Cassa et al.
- pLI scores from gnomAD v.2.1.1.

* Significant P-values are marked in bold and with an asterisk

| Genetic burden | Phenotype | Odds ratio (OR)/ effect size (ES) | P-value | Corrected p-value | Observations |
|---|---|---|---|---|---|
| Roulette | **childlessness** | OR = 2.152 | $1.10\times10^{-19}$ | **$1.10\times10^{-18}$*** | 374,671 |
| | **years of education** | ES = -2.820 | $2.17\times10^{-55}$ | **$2.17\times10^{-54}$*** | 372,949 |
| | **log(# of ICD-10 diagnoses)** | ES = 0.437 | $1.98\times10^{-34}$ | **$1.98\times10^{-33}$*** | 376,466 |
| | infertility | OR = 0.778 | 0.739 | 1 | 376,466 |
| | **living with a partner** | OR = 0.484 | $1.34\times10^{-20}$ | **$1.34\times10^{-19}$*** | 374,468 |
| | **fluid intelligence score** | ES = -1.430 | $2.33\times10^{-26}$ | **$2.33\times10^{-25}$*** | 120,725 |
| | hair color | OR = 0.831 | 0.115 | 1 | 375,729 |
| Weghorn | **childlessness** | OR = 2.060 | $1.51\times10^{-16}$ | **$1.51\times10^{-15}$*** | 374,704 |
| | **years of education** | ES = -2.583 | $7.78\times10^{-46}$ | **$7.78\times10^{-45}$*** | 372,982 |
| | log(# of ICD-10 diagnoses) | ES = 0.407 | $1.62\times10^{-29}$ | **$1.62\times10^{-28}$*** | 376,499 |
| | infertility | OR = 0.563 | 0.467 | 1 | 376,499 |
| | living with a partner | OR = 0.512 | $6.84\times10^{-17}$ | **$6.84\times10^{-16}$*** | 374,502 |
| | fluid intelligence score | ES = -1.183 | $1.83\times10^{-18}$ | **$1.83\times10^{-17}$*** | 120,730 |
| | hair color | OR = 0.922 | 0.486 | 1 | 375,762 |
| Cassa | **childlessness** | OR = 2.306 | $3.99\times10^{-18}$ | **$3.99\times10^{-17}$*** | 374,682 |
| | **years of education** | ES = -2.893 | $3.63\times10^{-46}$ | **$3.63\times10^{-45}$*** | 372,960 |
| | **log(# of ICD-10 diagnoses)** | ES = 0.480 | $9.39\times10^{-33}$ | **$9.39\times10^{-32}$*** | 376,477 |
| | infertility | OR = 0.649 | 0.615 | 1 | 376,477 |
| | **living with a partner** | OR = 0.472 | $2.73\times10^{-17}$ | **$2.73\times10^{-16}$*** | 374,480 |
| | **fluid intelligence score** | ES = -1.523 | $6.98\times10^{-24}$ | **$6.98\times10^{-23}$*** | 120,725 |
| | hair color | OR = 0.890 | 0.373 | 1 | 375,740 |
| pLI | **childlessness** | OR = 1.242 | $1.09\times10^{-16}$ | **$1.09\times10^{-15}$*** | 374,759 |
| | **years of education** | ES = -0.839 | $2.01\times10^{-54}$ | **$2.01\times10^{-53}$*** | 373,036 |
| | **log(# of ICD-10 diagnoses)** | ES = 0.127 | $1.36\times10^{-32}$ | **$1.36\times10^{-31}$*** | 376,554 |
| | infertility | OR = 0.899 | 0.637 | 1 | 376,554 |
| | **living with a partner** | OR = 0.818 | $3.64\times10^{-17}$ | **$3.64\times10^{-16}$*** | 374,557 |
| | **fluid intelligence score** | ES = -0.422 | $8.00\times10^{-26}$ | **$8.00\times10^{-25}$*** | 120,742 |
| | hair color | OR = 0.963 | 0.281 | 1 | 375,817 |

**Supplementary Table 6.** Regressions for various phenotypes performed on different disorder groups using the genetic burden based on $s_{het}$ scores from Roulette mutational model.

* Significant P-values are marked in bold and with an asterisk

| Gene panel | Phenotype | Odds ratio (OR)/ effect size (ES) | P-value | Corrected p-value | Observations |
|---|---|---|---|---|---|
| AR without ID | childlessness | OR = 1.422 | 0.067 | 0.944 | 369,265 |
| | years of education | ES = -1.096 | $4.34\times10^{-3}$ | 0.061 | 367,562 |
| | **log(# of ICD-10 diagnoses)** | ES = 0.261 | $6.15\times10^{-4}$ | **$8.61\times10^{-3}$*** | 371,041 |
| | fluid intelligence score | ES = -0.304 | 0.271 | 1 | 118,911 |
| | hair color | OR = 1.182 | 0.485 | 1 | 370,316 |
| **ID** | **childlessness** | OR = 2.756 | $1.69\times10^{-3}$ | **0.024*** | 373,755 |
| | **years of education** | ES = -3.931 | $2.03\times10^{-9}$ | **$2.84\times10^{-8}$*** | 372,032 |
| | **log(# of ICD-10 diagnoses)** | ES = 0.546 | $2.70\times10^{-5}$ | **$3.79\times10^{-4}$*** | 375,544 |
| | fluid intelligence score | ES = -1.420 | $5.50\times10^{-3}$ | 0.077 | 120,400 |
| | hair color | OR = 1.463 | 0.351 | 1 | 374,808 |
| Metabolic-ID | childlessness | OR = 1.825 | 0.232 | 1 | 374,775 |
| | years of education | ES = -0.230 | 0.818 | 1 | 373,052 |
| | log(# of ICD-10 diagnoses) | ES = 0.037 | 0.852 | 1 | 376,570 |
| | fluid intelligence score | ES = 0.582 | 0.432 | 1 | 120,746 |
| | hair color | OR = 0.274 | 0.047 | 0.656 | 375,833 |
| Blindness | childlessness | OR = 0.365 | 0.376 | 1 | 374,244 |
| | years of education | ES = -3.882 | 0.077 | 1 | 372,523 |
| | log(# of ICD-10 diagnoses) | ES = -0.004 | 0.993 | 1 | 376,036 |
| | fluid intelligence score | ES = -3.194 | 0.048 | 0.666 | 120,551 |
| | hair color | OR = 0.187 | 0.243 | 1 | 375,301 |
| Cilia+Kidney | childlessness | OR = 2.231 | 0.389 | 1 | 373,701 |
| | years of education | ES = -1.868 | 0.312 | 1 | 371,979 |
| | log(# of ICD-10 diagnoses) | ES = 0.108 | 0.769 | 1 | 375,492 |
| | fluid intelligence score | ES = -1.681 | 0.216 | 1 | 120,404 |
| | hair color | OR = 0.587 | 0.653 | 1 | 374,758 |
| Deafness | childlessness | OR = 7.133 | 0.175 | 1 | 374,504 |
| | years of education | ES = 3.087 | 0.299 | 1 | 372,784 |
| | log(# of ICD-10 diagnoses) | ES = -0.110 | 0.852 | 1 | 376,298 |
| | fluid intelligence score | ES = -1.353 | 0.569 | 1 | 120,660 |
| | hair color | OR = 0.901 | 0.955 | 1 | 375,562 |
| Dermatologic | childlessness | OR = 1.176 | 0.89 | 1 | 374,813 |
| | years of education | ES = -1.459 | 0.525 | 1 | 373,090 |
| | log(# of ICD-10 diagnoses) | ES = 0.328 | 0.472 | 1 | 376,608 |
| | fluid intelligence score | ES = 0.179 | 0.916 | 1 | 120,759 |
| | hair color | OR = 0.306 | 0.432 | 1 | 375,871 |

| | | | | | |
|---|---|---|---|---|---|
| Endocrine | childlessness | OR = 7.630 | 0.099 | 1 | 374,521 |
| | years of education | ES = 0.932 | 0.716 | 1 | 372,800 |
| | log(# of ICD-10 diagnoses) | ES = -0.743 | 0.145 | 1 | 376,316 |
| | fluid intelligence score | ES = -2.684 | 0.181 | 1 | 120,664 |
| | hair color | OR = 0.509 | 0.688 | 1 | 375,579 |
| Hematologic | childlessness | OR = 3.278 | 0.416 | 1 | 374,813 |
| | years of education | ES = 0.366 | 0.899 | 1 | 373,090 |
| | log(# of ICD-10 diagnoses) | ES = -0.084 | 0.884 | 1 | 376,608 |
| | fluid intelligence score | ES = -0.466 | 0.823 | 1 | 120,759 |
| | hair color | OR = 1.816 | 0.74 | 1 | 375,871 |
| Immune system | childlessness | OR = 1.290 | 0.549 | 1 | 374,762 |
| | years of education | ES = -1.998 | 0.023 | 0.319 | 373,039 |
| | log(# of ICD-10 diagnoses) | ES = 0.415 | 0.017 | 0.241 | 376,557 |
| | fluid intelligence score | ES = -0.206 | 0.722 | 1 | 120,743 |
| | hair color | OR = 2.228 | 0.095 | 1 | 375,821 |
| Neuromuscular | childlessness | OR = 0.618 | 0.446 | 1 | 373,960 |
| | years of education | ES = -2.503 | 0.039 | 0.541 | 372,245 |
| | log(# of ICD-10 diagnoses) | ES = 0.571 | 0.018 | 0.249 | 375,755 |
| | fluid intelligence score | ES = -1.912 | 0.03 | 0.426 | 120,499 |
| | hair color | OR = 1.938 | 0.37 | 1 | 375,021 |
| Skeletal | childlessness | OR = 0.735 | 0.73 | 1 | 374,789 |
| | years of education | ES = 0.465 | 0.786 | 1 | 373,067 |
| | log(# of ICD-10 diagnoses) | ES = -0.010 | 0.977 | 1 | 376,584 |
| | fluid intelligence score | ES = 1.430 | 0.245 | 1 | 120,751 |
| | hair color | OR = 2.204 | 0.439 | 1 | 375,847 |
| Metabolic | childlessness | OR = 1.580 | 0.283 | 1 | 374,212 |
| | years of education | ES = -2.463 | $4.16\times10^{-3}$ | 0.058 | 372,488 |
| | log(# of ICD-10 diagnoses) | ES = 0.175 | 0.303 | 1 | 376,004 |
| | fluid intelligence score | ES = -0.473 | 0.444 | 1 | 120,536 |
| | hair color | OR = 1.026 | 0.962 | 1 | 375,269 |
| Multi-system | childlessness | OR = 1.515 | 0.411 | 1 | 373,174 |
| | years of education | ES = 0.718 | 0.472 | 1 | 371,455 |
| | log(# of ICD-10 diagnoses) | ES = 0.444 | 0.024 | 0.342 | 374,961 |
| | fluid intelligence score | ES = 0.362 | 0.618 | 1 | 120,211 |
| | hair color | OR = 2.752 | 0.097 | 1 | 374,224 |

**Supplementary Table 7.** Regressions for various phenotypes performed on different disorder groups using the genetic burden based on $s_{het}$ scores from Weghorn et al.

* Significant P-values are marked in bold and with an asterisk

| Gene panel | Phenotype | Odds ratio (OR)/ effect size (ES) | P-value | Corrected p-value | Observations |
|---|---|---|---|---|---|
| AR without ID | childlessness | OR = 1.783 | $8.69\times10^{-3}$ | 0.122 | 357,686 |
| | years of education | ES = -0.564 | 0.199 | 1 | 356,016 |
| | log(# of ICD-10 diagnoses) | ES = 0.112 | 0.198 | 1 | 359,400 |
| | fluid intelligence score | ES = -0.373 | 0.249 | 1 | 115,115 |
| | hair color | OR = 1.465 | 0.16 | 1 | 358,700 |
| **ID** | childlessness | OR = 1.925 | 0.017 | 0.241 | 370,314 |
| | **years of education** | ES = -2.502 | $8.46\times10^{-6}$ | **$1.18\times10^{-4}$*** | 368,616 |
| | log(# of ICD-10 diagnoses) | ES = 0.133 | 0.232 | 1 | 372,086 |
| | fluid intelligence score | ES = -0.663 | 0.123 | 1 | 119,282 |
| | hair color | OR = 1.009 | 0.98 | 1 | 371,359 |
| Metabolic-ID | childlessness | OR = 1.579 | 0.346 | 1 | 372,390 |
| | years of education | ES = -0.593 | 0.537 | 1 | 370,670 |
| | log(# of ICD-10 diagnoses) | ES = 0.024 | 0.898 | 1 | 374,172 |
| | fluid intelligence score | ES = 0.476 | 0.519 | 1 | 119,928 |
| | hair color | OR = 0.403 | 0.153 | 1 | 373,441 |
| Blindness | childlessness | OR = 0.202 | 0.314 | 1 | 373,899 |
| | years of education | ES = -0.968 | 0.741 | 1 | 372,182 |
| | log(# of ICD-10 diagnoses) | ES = 0.515 | 0.377 | 1 | 375,691 |
| | fluid intelligence score | ES = -2.903 | 0.151 | 1 | 120,438 |
| | hair color | OR = 0.026 | 0.09 | 1 | 374,957 |
| Cilia+Kidney | childlessness | OR = 1.633 | 0.713 | 1 | 373,294 |
| | years of education | ES = 2.079 | 0.428 | 1 | 371,571 |
| | log(# of ICD-10 diagnoses) | ES = 0.037 | 0.944 | 1 | 375,082 |
| | fluid intelligence score | ES = -1.471 | 0.45 | 1 | 120,278 |
| | hair color | OR = 0.120 | 0.228 | 1 | 374,349 |
| Deafness | childlessness | OR = 11.911 | 0.02 | 0.276 | 372,833 |
| | years of education | ES = 2.873 | 0.209 | 1 | 371,117 |
| | log(# of ICD-10 diagnoses) | ES = -0.191 | 0.675 | 1 | 374,620 |
| | fluid intelligence score | ES = -1.191 | 0.498 | 1 | 120,094 |
| | hair color | OR = 2.571 | 0.484 | 1 | 373,887 |
| Dermatologic | childlessness | OR = 1.143 | 0.921 | 1 | 374,371 |
| | years of education | ES = -2.119 | 0.418 | 1 | 372,652 |
| | log(# of ICD-10 diagnoses) | ES = 0.542 | 0.299 | 1 | 376,166 |
| | fluid intelligence score | ES = 0.118 | 0.948 | 1 | 120,622 |
| | hair color | OR = 1.284 | 0.88 | 1 | 375,430 |

| | | | | | |
|---|---|---|---|---|---|
| Endocrine | childlessness | OR = 2.150 | 0.404 | 1 | 374,382 |
| | years of education | ES = 1.681 | 0.374 | 1 | 372,661 |
| | log(# of ICD-10 diagnoses) | ES = -0.489 | 0.19 | 1 | 376,175 |
| | fluid intelligence score | ES = -2.332 | 0.089 | 1 | 120,623 |
| | hair color | OR = 0.470 | 0.561 | 1 | 375,438 |
| Hematologic | childlessness | OR = 1.672 | 0.687 | 1 | 374,678 |
| | years of education | ES = -1.815 | 0.47 | 1 | 372,955 |
| | log(# of ICD-10 diagnoses) | ES = -0.007 | 0.99 | 1 | 376,473 |
| | fluid intelligence score | ES = -0.883 | 0.621 | 1 | 120,711 |
| | hair color | OR = 2.326 | 0.585 | 1 | 375,736 |
| Immune system | childlessness | OR = 1.432 | 0.557 | 1 | 373,936 |
| | years of education | ES = -1.723 | 0.159 | 1 | 372,216 |
| | log(# of ICD-10 diagnoses) | ES = 0.317 | 0.191 | 1 | 375,725 |
| | fluid intelligence score | ES = -0.713 | 0.403 | 1 | 120,477 |
| | hair color | OR = 1.508 | 0.585 | 1 | 374,990 |
| Neuromuscular | childlessness | OR = 0.750 | 0.785 | 1 | 373,295 |
| | years of education | ES = -5.641 | $5.70\times10^{-3}$ | 0.08 | 371,587 |
| | log(# of ICD-10 diagnoses) | ES = 0.830 | 0.041 | 0.572 | 375,087 |
| | fluid intelligence score | ES = -3.220 | 0.03 | 0.418 | 120,292 |
| | hair color | OR = 3.298 | 0.338 | 1 | 374,353 |
| Skeletal | childlessness | OR = 4.254 | 0.078 | 1 | 374,401 |
| | years of education | ES = -0.842 | 0.616 | 1 | 372,677 |
| | log(# of ICD-10 diagnoses) | ES = -0.016 | 0.962 | 1 | 376,190 |
| | fluid intelligence score | ES = 0.643 | 0.615 | 1 | 120,626 |
| | hair color | OR = 2.948 | 0.276 | 1 | 375,453 |
| Metabolic | childlessness | OR = 2.597 | 0.052 | 0.722 | 372,186 |
| | years of education | ES = -2.251 | 0.023 | 0.32 | 370,469 |
| | log(# of ICD-10 diagnoses) | ES = -0.023 | 0.905 | 1 | 373,969 |
| | fluid intelligence score | ES = -0.206 | 0.781 | 1 | 119,871 |
| | hair color | OR = 2.710 | 0.094 | 1 | 373,237 |
| Multi-system | childlessness | OR = 1.383 | 0.522 | 1 | 371,004 |
| | years of education | ES = 1.555 | 0.119 | 1 | 369,296 |
| | log(# of ICD-10 diagnoses) | ES = 0.355 | 0.073 | 1 | 372,781 |
| | fluid intelligence score | ES = 0.587 | 0.42 | 1 | 119,499 |
| | hair color | OR = 2.281 | 0.176 | 1 | 372,053 |

**Supplementary Table 8.** Regressions for various phenotypes performed on different disorder groups using the genetic burden based on $s_{het}$ scores from Cassa et al.

* Significant P-values are marked in bold and with an asterisk

| Gene panel | Phenotype | Odds ratio (OR)/ effect size (ES) | P-value | Corrected p-value | Observations |
|---|---|---|---|---|---|
| AR without ID | childlessness | OR = 1.664 | 0.054 | 0.755 | 357,033 |
| | years of education | ES = -0.817 | 0.122 | 1 | 355,367 |
| | log(# of ICD-10 diagnoses) | ES = 0.258 | 0.014 | 0.193 | 358,742 |
| | fluid intelligence score | ES = -0.373 | 0.342 | 1 | 114,900 |
| | hair color | OR = 1.499 | 0.213 | 1 | 358,043 |
| **ID** | childlessness | OR = 2.448 | $8.17\times10^{-3}$ | 0.114 | 370,816 |
| | **years of education** | ES = -3.146 | $6.51\times10^{-6}$ | **$9.11\times10^{-5}$*** | 369,117 |
| | log(# of ICD-10 diagnoses) | ES = 0.288 | 0.037 | 0.524 | 372,590 |
| | fluid intelligence score | ES = -1.014 | 0.054 | 0.751 | 119,436 |
| | hair color | OR = 0.824 | 0.665 | 1 | 371,864 |
| Metabolic-ID | childlessness | OR = 2.027 | 0.265 | 1 | 372,983 |
| | years of education | ES = 0.293 | 0.817 | 1 | 371,260 |
| | log(# of ICD-10 diagnoses) | ES = 0.070 | 0.782 | 1 | 374,767 |
| | fluid intelligence score | ES = 0.755 | 0.445 | 1 | 120,139 |
| | hair color | OR = 0.154 | 0.03 | 0.418 | 374,036 |
| Blindness | childlessness | OR = 0.121 | 0.132 | 1 | 373,899 |
| | years of education | ES = 0.540 | 0.82 | 1 | 372,182 |
| | log(# of ICD-10 diagnoses) | ES = 0.905 | 0.056 | 0.785 | 375,691 |
| | fluid intelligence score | ES = -1.377 | 0.435 | 1 | 120,438 |
| | hair color | OR = 0.093 | 0.178 | 1 | 374,957 |
| Cilia+Kidney | childlessness | OR = 0.823 | 0.886 | 1 | 372,972 |
| | years of education | ES = -1.578 | 0.542 | 1 | 371,250 |
| | log(# of ICD-10 diagnoses) | ES = 0.712 | 0.163 | 1 | 374,757 |
| | fluid intelligence score | ES = -1.265 | 0.537 | 1 | 120,154 |
| | hair color | OR = 0.995 | 0.998 | 1 | 374,025 |
| Deafness | childlessness | OR = 22.556 | 0.08 | 1 | 372,793 |
| | years of education | ES = 2.374 | 0.523 | 1 | 371,077 |
| | log(# of ICD-10 diagnoses) | ES = -0.423 | 0.567 | 1 | 374,580 |
| | fluid intelligence score | ES = -3.645 | 0.241 | 1 | 120,078 |
| | hair color | OR = 2.728 | 0.654 | 1 | 373,847 |
| Dermatologic | childlessness | OR = 1.529 | 0.84 | 1 | 374,120 |
| | years of education | ES = -2.049 | 0.618 | 1 | 372,403 |
| | log(# of ICD-10 diagnoses) | ES = 0.522 | 0.524 | 1 | 375,915 |
| | fluid intelligence score | ES = -0.063 | 0.983 | 1 | 120,540 |
| | hair color | OR = 1.713 | 0.835 | 1 | 375,180 |

| | | | | | |
|---|---|---|---|---|---|
| Endocrine | childlessness | OR = 12.983 | 0.157 | 1 | 374,329 |
| | years of education | ES = 1.414 | 0.71 | 1 | 372,607 |
| | log(# of ICD-10 diagnoses) | ES = -1.195 | 0.112 | 1 | 376,121 |
| | fluid intelligence score | ES = -4.448 | 0.129 | 1 | 120,607 |
| | hair color | OR = 0.054 | 0.288 | 1 | 375,384 |
| Hematologic | childlessness | OR = 30.706 | 0.207 | 1 | 374,520 |
| | years of education | ES = 2.979 | 0.581 | 1 | 372,797 |
| | log(# of ICD-10 diagnoses) | ES = -0.092 | 0.932 | 1 | 376,314 |
| | fluid intelligence score | ES = 0.626 | 0.872 | 1 | 120,659 |
| | hair color | OR = 1.228 | 0.952 | 1 | 375,577 |
| Immune system | childlessness | OR = 1.545 | 0.513 | 1 | 373,936 |
| | years of education | ES = -1.292 | 0.339 | 1 | 372,216 |
| | log(# of ICD-10 diagnoses) | ES = 0.562 | 0.036 | 0.509 | 375,725 |
| | fluid intelligence score | ES = -0.441 | 0.636 | 1 | 120,477 |
| | hair color | OR = 1.369 | 0.707 | 1 | 374,990 |
| Neuromuscular | childlessness | OR = 1.180 | 0.846 | 1 | 373,295 |
| | years of education | ES = -3.890 | 0.021 | 0.297 | 371,587 |
| | log(# of ICD-10 diagnoses) | ES = 0.540 | 0.108 | 1 | 375,087 |
| | fluid intelligence score | ES = -2.624 | 0.029 | 0.401 | 120,292 |
| | hair color | OR = 2.183 | 0.443 | 1 | 374,353 |
| Skeletal | childlessness | OR = 1.511 | 0.692 | 1 | 374,401 |
| | years of education | ES = -0.202 | 0.921 | 1 | 372,677 |
| | log(# of ICD-10 diagnoses) | ES = 0.139 | 0.731 | 1 | 376,190 |
| | fluid intelligence score | ES = 0.975 | 0.504 | 1 | 120,626 |
| | hair color | OR = 5.716 | 0.128 | 1 | 375,453 |
| Metabolic | childlessness | OR = 1.651 | 0.364 | 1 | 372,090 |
| | years of education | ES = -3.220 | $3.59\times10^{-3}$ | 0.05 | 370,373 |
| | log(# of ICD-10 diagnoses) | ES = 0.058 | 0.792 | 1 | 373,871 |
| | fluid intelligence score | ES = -0.333 | 0.687 | 1 | 119,845 |
| | hair color | OR = 3.753 | 0.041 | 0.578 | 373,138 |
| Multi-system | childlessness | OR = 2.019 | 0.286 | 1 | 370,875 |
| | years of education | ES = 2.303 | 0.081 | 1 | 369,167 |
| | log(# of ICD-10 diagnoses) | ES = 0.553 | 0.034 | 0.481 | 372,651 |
| | fluid intelligence score | ES = 0.672 | 0.512 | 1 | 119,443 |
| | hair color | OR = 3.621 | 0.1 | 1 | 371,923 |

**Supplementary Table 9.** Regressions for various phenotypes performed on different disorder groups using the genetic burden based on pLI.

* Significant P-values are marked in bold and with an asterisk

| Gene panel | Phenotype | Odds ratio (OR)/ effect size (ES) | P-value | Corrected p-value | Observations |
|---|---|---|---|---|---|
| AR without ID | childlessness | OR = 1.103 | 0.021 | 0.289 | 344,184 |
| | **years of education** | ES = -0.349 | $4.03\times10^{-5}$ | **$5.65\times10^{-4}$*** | 342,569 |
| | **log(# of ICD-10 diagnoses)** | ES = 0.058 | $6.47\times10^{-4}$ | **$9.06\times10^{-3}$*** | 345,837 |
| | fluid intelligence score | ES = -0.117 | 0.062 | 0.861 | 110,697 |
| | hair color | OR = 1.037 | 0.491 | 1 | 345,147 |
| **ID** | **childlessness** | OR = 1.219 | $1.69\times10^{-3}$ | **0.024*** | 362,813 |
| | **years of education** | ES = -0.512 | $7.73\times10^{-5}$ | **$1.08\times10^{-3}$*** | 361,143 |
| | **log(# of ICD-10 diagnoses)** | ES = 0.108 | $2.54\times10^{-5}$ | **$3.55\times10^{-4}$*** | 364,553 |
| | fluid intelligence score | ES = -0.249 | 0.014 | 0.194 | 116,841 |
| | hair color | OR = 1.060 | 0.469 | 1 | 363,838 |
| **Metabolic-ID** | childlessness | OR = 0.983 | 0.881 | 1 | 370,560 |
| | **years of education** | ES = -0.626 | $4.92\times10^{-3}$ | 0.069 | 368,852 |
| | log(# of ICD-10 diagnoses) | ES = 0.054 | 0.221 | 1 | 372,331 |
| | fluid intelligence score | ES = -0.056 | 0.746 | 1 | 119,292 |
| | hair color | OR = 0.672 | 0.014 | 0.195 | 371,597 |
| Blindness | childlessness | OR = 1.003 | 0.991 | 1 | 370,347 |
| | years of education | ES = -1.130 | 0.028 | 0.393 | 368,646 |
| | log(# of ICD-10 diagnoses) | ES = 0.209 | 0.04 | 0.561 | 372,120 |
| | fluid intelligence score | ES = -0.515 | 0.183 | 1 | 119,322 |
| | hair color | OR = 0.700 | 0.331 | 1 | 371,391 |
| Cilia+Kidney | childlessness | OR = 1.167 | 0.425 | 1 | 371,301 |
| | years of education | ES = -0.570 | 0.144 | 1 | 369,580 |
| | log(# of ICD-10 diagnoses) | ES = 0.077 | 0.322 | 1 | 373,073 |
| | fluid intelligence score | ES = -0.209 | 0.464 | 1 | 119,597 |
| | hair color | OR = 1.284 | 0.266 | 1 | 372,343 |
| Deafness | childlessness | OR = 1.771 | 0.052 | 0.73 | 372,981 |
| | years of education | ES = 0.114 | 0.86 | 1 | 371,263 |
| | log(# of ICD-10 diagnoses) | ES = 0.104 | 0.415 | 1 | 374,768 |
| | fluid intelligence score | ES = -0.962 | 0.101 | 1 | 120,144 |
| | hair color | OR = 1.360 | 0.398 | 1 | 374,034 |
| Dermatologic | childlessness | OR = 1.119 | 0.57 | 1 | 373,295 |
| | years of education | ES = -0.446 | 0.258 | 1 | 371,578 |
| | log(# of ICD-10 diagnoses) | ES = 0.200 | 0.011 | 0.154 | 375,086 |
| | fluid intelligence score | ES = 0.199 | 0.484 | 1 | 120,245 |
| | hair color | OR = 0.791 | 0.381 | 1 | 374,352 |

| | | | | | |
|---|---|---|---|---|---|
| Endocrine | childlessness | OR = 1.740 | 0.015 | 0.209 | 374,035 |
| | years of education | ES = 0.065 | 0.896 | 1 | 372,316 |
| | log(# of ICD-10 diagnoses) | ES = -0.222 | 0.025 | 0.343 | 375,828 |
| | fluid intelligence score | ES = -0.793 | 0.043 | 0.598 | 120,480 |
| | hair color | OR = 0.709 | 0.334 | 1 | 375,091 |
| Hematologic | childlessness | OR = 0.594 | 0.489 | 1 | 374,305 |
| | years of education | ES = -0.544 | 0.691 | 1 | 372,587 |
| | log(# of ICD-10 diagnoses) | ES = 0.098 | 0.719 | 1 | 376,099 |
| | fluid intelligence score | ES = -0.113 | 0.905 | 1 | 120,594 |
| | hair color | OR = 0.214 | 0.19 | 1 | 375,364 |
| Immune system | childlessness | OR = 1.078 | 0.527 | 1 | 373,850 |
| | years of education | ES = -0.143 | 0.546 | 1 | 372,128 |
| | log(# of ICD-10 diagnoses) | ES = 0.112 | 0.018 | 0.248 | 375,638 |
| | fluid intelligence score | ES = -0.270 | 0.12 | 1 | 120,444 |
| | hair color | OR = 1.306 | 0.05 | 0.696 | 374,903 |
| **Neuromuscular** | childlessness | OR = 0.977 | 0.834 | 1 | 372,033 |
| | **years of education** | ES = -0.714 | $7.72\times10^{-4}$ | **0.011*** | 370,330 |
| | log(# of ICD-10 diagnoses) | ES = 0.100 | 0.017 | 0.244 | 373,824 |
| | fluid intelligence score | ES = -0.257 | 0.093 | 1 | 119,872 |
| | hair color | OR = 0.959 | 0.758 | 1 | 373,093 |
| Skeletal | childlessness | OR = 1.146 | 0.509 | 1 | 374,155 |
| | years of education | ES = 0.168 | 0.685 | 1 | 372,428 |
| | log(# of ICD-10 diagnoses) | ES = -0.030 | 0.709 | 1 | 375,942 |
| | fluid intelligence score | ES = 0.647 | 0.038 | 0.531 | 120,519 |
| | hair color | OR = 1.014 | 0.958 | 1 | 375,205 |
| Metabolic | childlessness | OR = 1.149 | 0.168 | 1 | 368,658 |
| | years of education | ES = -0.396 | 0.052 | 0.723 | 366,944 |
| | log(# of ICD-10 diagnoses) | ES = 0.038 | 0.351 | 1 | 370,420 |
| | fluid intelligence score | ES = -0.063 | 0.667 | 1 | 118,711 |
| | hair color | OR = 1.166 | 0.211 | 1 | 369,692 |
| Multi-system | childlessness | OR = 1.164 | 0.108 | 1 | 363,877 |
| | years of education | ES = 0.010 | 0.958 | 1 | 362,203 |
| | log(# of ICD-10 diagnoses) | ES = -0.006 | 0.881 | 1 | 365,624 |
| | fluid intelligence score | ES = -0.012 | 0.933 | 1 | 117,202 |
| | hair color | OR = 1.210 | 0.095 | 1 | 364,907 |

**Supplementary Table 10.** Regressions performed for males and females on PLPs in recessive genes using several sources for genetic burden scores:

- Roulette-based scores from Seplyarskiy et al.
- Weghorn et al. using a demographic model that includes genetic drift
- Cassa et al.
- pLI scores from gnomAD v.2.1.1.

* Significant P-values are marked in bold and with an asterisk

| Genetic burden | Gene set | Gender | Phenotype | Odds ratio (OR)/ effect size (ES) | P-value | Corrected p-value | Observations |
|---|---|---|---|---|---|---|---|
| Roulette | 1,929 recessive | male | **childlessness** | OR = 1.997 | $3.05\times10^{-3}$ | **0.037*** | 169,906 |
| | | | **years of education** | ES = -1.693 | $5.38\times10^{-4}$ | **$6.45\times10^{-3}$*** | 169,798 |
| | | | hair color | OR = 1.592 | 0.141 | 1 | 170,888 |
| | | female | childlessness | OR = 1.436 | 0.126 | 1 | 198,318 |
| | | | **years of education** | ES = -1.990 | $1.06\times10^{-5}$ | **$1.27\times10^{-4}$*** | 196,722 |
| | | | hair color | OR = 1.050 | 0.86 | 1 | 198,382 |
| Weghorn | 1,929 recessive | male | childlessness | OR = 1.822 | 0.016 | 0.192 | 163,089 |
| | | | years of education | ES = -1.144 | 0.027 | 0.325 | 162,973 |
| | | | hair color | OR = 2.283 | 0.011 | 0.128 | 164,028 |
| | | female | childlessness | OR = 1.811 | 0.016 | 0.192 | 190,336 |
| | | | years of education | ES = -1.321 | $5.51\times10^{-3}$ | 0.066 | 188,804 |
| | | | hair color | OR = 0.796 | 0.442 | 1 | 190,396 |
| Cassa | 1,929 recessive | male | childlessness | OR = 2.345 | $4.65\times10^{-3}$ | 0.056 | 163,018 |
| | | | years of education | ES = -1.661 | $8.87\times10^{-3}$ | 0.106 | 162,902 |
| | | | hair color | OR = 1.978 | 0.088 | 1 | 163,959 |
| | | female | childlessness | OR = 1.709 | 0.072 | 0.864 | 190,228 |
| | | | years of education | ES = -1.513 | $8.24\times10^{-3}$ | 0.099 | 188,699 |
| | | | hair color | OR = 0.836 | 0.615 | 1 | 190,286 |
| pLI | 1,929 recessive | male | **childlessness** | OR = 1.237 | $1.90\times10^{-5}$ | **$2.28\times10^{-4}$*** | 155,751 |
| | | | **years of education** | ES = -0.394 | $1.78\times10^{-4}$ | **$2.13\times10^{-3}$*** | 155,631 |
| | | | hair color | OR = 1.106 | 0.135 | 1 | 156,637 |
| | | female | childlessness | OR = 1.041 | 0.436 | 1 | 181,728 |
| | | | **years of education** | ES = -0.415 | $1.76\times10^{-5}$ | **$2.11\times10^{-4}$*** | 180,261 |
| | | | hair color | OR = 1.006 | 0.922 | 1 | 181,786 |

**Supplementary Table 11.** Regressions performed for males and females on singleton LoF variants in non-recessive highly constrained ($s_{het} \geq 0.15$) genes using several sources for genetic burden scores:

- Roulette-based scores from Seplyarskiy et al.
- Weghorn et al. using a demographic model that includes genetic drift
- Cassa et al.
- pLI scores from gnomAD v.2.1.1.

* Significant P-values are marked in bold and with an asterisk

| Genetic burden | Gene set | Gender | Phenotype | Odds ratio (OR)/ effect size (ES) | P-value | Corrected p-value | Observations |
|---|---|---|---|---|---|---|---|
| Roulette | Non-recessive highly constrained | male | **childlessness** | OR = 2.919 | $3.51\times10^{-20}$ | **$4.22\times10^{-19}$*** | 172,815 |
| | | | **years of education** | ES = -3.096 | $2.16\times10^{-32}$ | **$2.59\times10^{-31}$*** | 172,711 |
| | | | hair color | OR = 0.960 | 0.817 | 1 | 173,810 |
| | | female | **childlessness** | OR = 1.532 | $5.90\times10^{-4}$ | **$7.07\times10^{-3}$*** | 201,856 |
| | | | **years of education** | ES = -2.642 | $5.63\times10^{-27}$ | **$6.76\times10^{-26}$*** | 200,238 |
| | | | hair color | OR = 0.765 | 0.088 | 1 | 201,919 |
| Weghorn | Non-recessive highly constrained | male | **childlessness** | OR = 2.850 | $4.76\times10^{-18}$ | **$5.71\times10^{-17}$*** | 172,836 |
| | | | **years of education** | ES = -2.918 | $2.40\times10^{-28}$ | **$2.87\times10^{-27}$*** | 172,731 |
| | | | hair color | OR = 1.126 | 0.498 | 1 | 173,831 |
| | | female | childlessness | OR = 1.436 | $4.64\times10^{-3}$ | 0.056 | 201,868 |
| | | | **years of education** | ES = -2.351 | $2.44\times10^{-21}$ | **$2.92\times10^{-20}$*** | 200,251 |
| | | | hair color | OR = 0.812 | 0.182 | 1 | 201,931 |
| Cassa | Non-recessive highly constrained | male | **childlessness** | OR = 3.261 | $1.83\times10^{-19}$ | **$2.20\times10^{-18}$*** | 172,824 |
| | | | **years of education** | ES = -3.016 | $3.48\times10^{-25}$ | **$4.18\times10^{-24}$*** | 172,719 |
| | | | hair color | OR = 1.083 | 0.68 | 1 | 173,819 |
| | | female | **childlessness** | OR = 1.512 | $3.88\times10^{-3}$ | **0.047*** | 201,858 |
| | | | **years of education** | ES = -2.905 | $3.59\times10^{-25}$ | **$4.30\times10^{-24}$*** | 200,241 |
| | | | hair color | OR = 0.794 | 0.193 | 1 | 201,921 |
| pLI | Non-recessive highly constrained | male | **childlessness** | OR = 1.380 | $5.86\times10^{-19}$ | **$7.03\times10^{-18}$*** | 172,872 |
| | | | **years of education** | ES = -0.926 | $5.75\times10^{-32}$ | **$6.90\times10^{-31}$*** | 172,767 |
| | | | hair color | OR = 0.999 | 0.988 | 1 | 173,867 |
| | | female | childlessness | OR = 1.107 | $7.57\times10^{-3}$ | 0.091 | 201,887 |
| | | | **years of education** | ES = -0.784 | $1.46\times10^{-26}$ | **$1.75\times10^{-25}$*** | 200,269 |
| | | | hair color | OR = 0.945 | 0.221 | 1 | 201,950 |

**Supplementary Table 12**. Permutation test for CR scores for 13 disorder groups on 3 European cohorts (**Methods**): the UK Biobank, Dutch and Estonian.

Scores for the Dutch and Estonian cohorts are taken from Fridman et al.[9]
* Significant P-values are marked in bold and with an asterisk

| Panel | UK biobank (n=378,751) | | Dutch cohort (n=4,120) | | Estonian cohort (n=2,327) | |
|---|---|---|---|---|---|---|
| | CR | corrected p-value | CR | corrected p-value | CR | corrected p-value |
| Hematologic | 123.95 | 0.221 | 19.1 | 1.00 | 30.1 | 1.00 |
| Neuromuscular | 77.74 | 0.296 | 47.3 | 1.00 | 9.5 | 1.00 |
| **ID** | 66.86 | **0.005*** | 48.0 | **0.01*** | 40.9 | **0.04*** |
| Skeletal | 60.24 | 1.0000 | 123.0 | **0.01*** | 91.9 | 0.29 |
| Cilia+Kidney | 41.42 | 1.0000 | 36.7 | 1.00 | 18.2 | 1.00 |
| Dermatologic | 33.31 | 1.0000 | 29.1 | 1.00 | 22.8 | 1.00 |
| Metabolic-ID | 33.36 | 1.0000 | 14.2 | 1.00 | 20.1 | 1.00 |
| Multi-system | 28.81 | 1.0000 | 18.7 | 1.00 | 32.6 | 1.00 |
| Blindness | 28.56 | 1.0000 | 7.1 | 1.00 | 6.6 | 1.00 |
| Immune system | 20.99 | 1.0000 | 22.6 | 1.00 | 28.7 | 1.00 |
| Endocrine | 18.60 | 1.0000 | 14.4 | 1.00 | 13.4 | 1.00 |
| Deafness | 15.80 | 1.0000 | 21.7 | 1.00 | 6.6 | 1.00 |
| Metabolic | 9.40 | 1.0000 | 9.5 | 1.00 | 11.3 | 1.00 |

**Supplementary Table 13**. $s_{het}$ (from Roulette) distribution across disorder groups for recessive genes.

| Disorder group | # of genes | # of genes with $s_{het}$ score | # of genes with higher $s_{het}$ score (>=0.1) |
|---|---|---|---|
| All genes | 1,929 | 1,890 (97.9%) | 125 (6.6%) |
| ID | 363 | 357 (98.3%) | 37 (10.4%) |
| Metabolic-ID | 353 | 352 (99.7%) | 10 (2.8%) |
| Metabolic | 318 | 314 (98.7%) | 13 (4.1%) |
| Multi-system | 271 | 263 (97.0%) | 13 (4.9%) |
| Neuromuscular | 113 | 109 (96.5%) | 11 (10.1%) |
| Immune system | 97 | 87 (95.6%) | 18 (20.7%) |
| Blindness | 85 | 80 (94.1%) | 2 (2.5%) |
| Cilia+Kidney | 74 | 72 (97.3%) | 5 (6.9%) |
| Skeletal | 61 | 60 (98.4%) | 7 (11.7%) |
| Dermatologic | 59 | 59 (100.0%) | 1 (1.7%) |
| Hematologic | 39 | 39 (100.0%) | 1 (2.6%) |
| Deafness | 38 | 36 (94.7%) | 2 (5.6%) |
| Endocrine | 33 | 32 (97.0%) | 3 (9.4%) |

**Supplementary Table 14.** Description of the phenotypes used from the UK Biobank.

| Phenotype | UKBB field number | Notes |
|---|---|---|
| Sex | 31 | Used to determine sex. |
| Age at recruitment | 21022 | Used to determine age. |
| Number of children fathered | 2405 | Individuals that replied with a value of -1 (“Do not know”) or -3 (“Prefer not to answer”) or >7 were assigned N/A value |
| Number of live births | 2734 | Individuals that replied with a value of -1 (“Do not know”) or -3 (“Prefer not to answer”) or >7 were assigned N/A value |
| Qualifications | 6138 | This field was used to determine educational attainment (in years of education, Methods). The highest qualification reported was mapped to the International Standard Classification for Education (ISCED) coding for years of education:<br>*College or University degree* - 20 years<br>*NVQ or HND or HNC or equivalent* - 19 years<br>*Other prof. qual. eg: nursing, teaching* - 15 years<br>*A levels/AS levels or equivalent* - 13 years<br>*O levels/GCSEs or equivalent* - 10 years<br>*CSEs or equivalent* - 10 years<br>*None of the above* - 7 years<br>We excluded participants opted for “Prefer not to answer” from the relevant analyses. |
| Fluid intelligence score | 20016 | Fluid intelligence score measures the capacity to solve problems that require logic and reasoning ability, independent of acquired knowledge.<br><br>It’s calculated as an unweighted sum of the number of correct answers given to the 13 fluid intelligence questions. |
| Hair color | 1747 | Individuals that replied 1 (“Blonde”) were assigned a value of 1, individuals that replied -1 (“Do not know”) or -3 (“Prefer not to answer”) were assigned N/A value, all other were assigned a value of 1 |
| Genetic ethnic grouping | 22006 | Used to determine ancestry as described in the Methods. |
| Genetic principal components | 22009 | Used for the regression analyses as described in the Methods. |
| Diagnoses - main ICD10 | 41202 | This field contains a summary of the distinct primary/main diagnosis codes a participant has recorded across all their hospital inpatient records. Used to determine the total number of main diagnoses. |
| Diagnoses - secondary ICD10 | 41204 | This field contains a summary of the distinct secondary diagnosis codes a participant has recorded across all their hospital inpatient records. Used to determine the total number of secondary diagnoses.<br><br>Together with the main ICD10 diagnoses field determines the total number of diagnoses. |
| Infertility | 41202, 41204 | This phenotype is based on main and secondary ICD10 diagnoses. Male infertility code: N46, female infertility code: N97. |
| Indices of | 76 | Indices of Multiple Deprivation come from a UK government qualitative study |

| | | |
|---|---|---|
| multiple deprivation | | of deprived areas in British local councils. For the analyses on deprivation, we only used the samples for whom England scores were available in order to maintain consistency between scores. We included the following data:<br>- Index of multiple deprivation<br>- Education score<br>- Health score<br>- Housing score<br>- Income score |

**Supplementary table 15.** Overview of the analyses and Bonferroni correction.

| Analysis | Variants set | Gene set | Samples set | Phenotype | Predictor | Bonferroni correction |
|---|---|---|---|---|---|---|
| Selection / fitness effect analysis | PLPs | 1,929 AR | all | childlessness | # of PLPs | correction was applied for 10 tests in total<br><br>significance level is **0.005** |
| | | | | childlessness | carriership status | |
| | | | | childlessness | $s_{het}$ | |
| | | | | hair color | | |
| | | | | childlessness | $s_{het}$ [PLPs] + $s_{het}$ [LoFs] | |
| | | | | hair color | | |
| | | | all without LoF carriers | childlessness | $s_{het}$ | |
| | | | | hair color | | |
| | LoFs | non-AR s-het>=0.15 | all | childlessness | $s_{het}$ | |
| | | | | hair color | | |
| Phenotypic analysis | PLPs<br>———<br>LoFs | 1,929 AR<br>———<br>non-AR s-het>=0.15 | all | years of education | $s_{het}$ | correction was applied for 10 tests in total<br><br>significance level is **0.005** |
| | | | | # ICD-10 diagnoses | | |
| | | | | infertility | | |
| | | | | living with a partner | | |
| | | | | fluid intelligence | | |
| Disorder groups analysis | PLPs | 13 disorder groups + all non-ID genes combined | all | childlessness | $s_{het}$ | correction was applied for 14 tests in total<br><br>significance level per phenotype is **$3.57\times10^{-3}$** |
| | | | | years of education | | |
| | | | | # ICD-10 diagnoses | | |
| | | | | fluid intelligence | | |
| | | | | hair color | | |
| Sex-specific differencies | PLPs<br>———<br>LoFs | 1,929 AR<br>———<br>non-AR s-het>=0.15 | males<br>———<br>females | childlessness | $s_{het}$ | correction was applied for 12 tests in total<br><br>significance level is **0.0042** |
| | | | | years of education | | |
| | | | | hair color | | |

**Supplementary Table 16**. Samples distribution for the number of variants per sample.

| Number of variants per sample | Number of samples for PLPs in recessive genes | Number of samples for rare (<=20 hets) synonymous variants in recessive genes |
|---|---|---|
| 1 | 108,179 | 78,437 |
| 2 | 106,539 | 95,286 |
| 3 | 68,639 | 79,915 |
| 4 | 33,546 | 52,553 |
| 5 | 12,829 | 28,724 |
| 6 | 4,179 | 13,661 |
| 7 | 1,152 | 5,763 |
| 8 | 242 | 2,165 |
| 9 | 57 | 787 |
| 10 | 11 | 254 |
| 11 | 1 | 75 |
| 12 | - | 24 |
| 13 | - | 7 |
| 14 | - | 5 |
